# Structural basis of receptor retro-translocation in peroxisomal protein import

**DOI:** 10.64898/2026.03.18.712702

**Authors:** Nathaniel W.M. Dempsey, Laurie Wang, Ningjian Gao, Kangting Zhao, Jessica Cope, Eunyong Park

## Abstract

Peroxisomes import all matrix proteins post-translationally from the cytosol, a process that requires recycling of cargo receptors across the peroxisomal membrane. The membrane-embedded ubiquitin ligase, composed of Pex2, Pex10, and Pex12, is central to this process, but its mechanism remains unclear. Here we determined cryo-electron microscopy structures of the *Saccharomyces cerevisiae* Pex2-10-12 complex in closed and open states bound to Pex8, an essential factor of previously undefined function. The structures reveal how Pex2-10-12 gates its retro-translocation pore to control receptor entry and how the closed-to-open transition repositions the Pex10 RING domain to enable receptor mono-ubiquitination. Pex8 docks onto Pex2-10-12 from the matrix and guides receptors into the pore. Functional analyses show that the receptor’s N-terminal segment downstream of its mono-ubiquitination site initiates a loop insertion into the pore. These findings establish how Pex2-10-12 coordinates receptor recognition, retro-translocation, and ubiquitination, providing the molecular basis for receptor recycling in peroxisomal protein import.

## INTRODUCTION

Peroxisomes are small single-membrane-bound organelles found in most eukaryotic cells. They perform various metabolic reactions, such as β-oxidation of fatty acids, detoxification of hydrogen peroxide, synthesis of plasmalogens, and the production and breakdown of reactive oxygen species. All lumenal (matrix) proteins of the peroxisome are imported from the cytosol by concerted actions of conserved proteins called peroxins (Pex proteins; for recent reviews, see ref. ^1,2^). These include cargo import receptors, docking and translocation machinery, and a receptor recycling system, disruptions of any of which impair import of matrix proteins. In humans, loss-of-function mutations in *PEX* genes often cause peroxisome biogenesis disorders, such as Zellweger spectrum disorders^3,4^. While significant progress has been made in understanding the functions of Pex proteins, much about their coordination and dynamics remains to be elucidated.

The peroxisomal import process begins with the recognition of cargo proteins by a soluble import receptor in the cytosol. Most cargo proteins contain a highly conserved C-terminal serine-lysine-leucine (SKL) tripeptide motif, or a variant (e.g., AKL and SHL), collectively known as a type-1 peroxisomal targeting signal (PTS1)^5^. Some matrix proteins utilize a type-2 peroxisomal targeting signal (PTS2), a nine-amino-acid sequence with the consensus R-(L/V/I)-XXXXX-(H/Q)-(L/A) (‘X’ represents positions with low conservation) typically located near the N-terminus^6^. PTS1 is recognized directly by the import receptor Pex5 through its C-terminal tetratricopeptide repeat (TPR) domain^7,8^, while PTS2 binds to the receptor—Pex5 in humans, or Pex18/Pex21 in yeast, which lack a TPR domain—via the adaptor protein Pex7 (ref. ^9–12^).

Upon cargo recognition, the import receptor transports the bound cargo across the peroxisomal membrane through an import pore complex composed of Pex13 and Pex14, and other associated proteins (e.g., Pex17 in yeast). Unlike protein translocation into the endoplasmic reticulum (ER) or mitochondria, peroxisomal import allows translocation of folded cargo proteins^13^. Recent studies have shown that the conduit is formed by Pex13, whose intrinsically disordered regions (IDRs) create a diffusion barrier by phase separation, analogous to that of the nuclear pore^14,15^. Pex5 contains a disordered N-terminal segment with affinity for Pex13’s IDRs and docking components like Pex14, enabling selective translocation of Pex5 and its bound cargo in a manner reminiscent of karyopherin-mediated nuclear import^16,17^. Fungal PTS2 receptors Pex18 and Pex21, which also possess an analogous disordered segment, use a similar mechanism but additionally rely on a Pex7–Pex13 interaction alongside the Pex18/21–Pex13 interaction for peroxisomal entry^11^.

Once imported into the peroxisomal matrix, import receptors must release their cargo and return to the cytosol for the next round of import. This recycling process is mediated by an E3 ubiquitin ligase complex embedded in the peroxisomal membrane, composed of Pex2, Pex10, and Pex12 (referred to as the Pex2-10-12 complex)^18–23^. The complex engages cargo-bound import receptors and catalyzes the mono-ubiquitination of a conserved cysteine residue near the N-terminus of the receptor^24^. Each subunit of the Pex2-10-12 complex contains a cytosolic RING finger (RF) domain, which together assemble into a heterotrimeric RING domain (hereafter ‘RING domain’ with ‘RF’ referring to individual subunits)^22,23^. This RING domain recruits ubiquitin and an E2 ubiquitin-conjugating enzyme—Pex4 in yeast and UbcH5a/b/c in humans—to catalyze receptor ubiquitination^25,26^.

The key step of import receptor recycling is retro-translocation of the receptors from the peroxisomal matrix back to the cytosol^27^. Studies have shown that the N-terminal region of matrix-imported Pex5 is exposed to the cytosol for ubiquitination^28,29^. A recent cryo-electron microscopy (cryo-EM) structure of the Pex2-10-12 complex from the thermophilic fungus *Thermothelomyces thermophilus* exhibited a pore of approximately 10 Å diameter within its transmembrane domain (TMD)^23^. Together, these data suggest that the Pex2-10-12 complex serves as a retro-translocation channel for receptor recycling.

Once mono-ubiquitinated, the receptors are recognized by the Pex1–Pex6 complex, a heterohexameric AAA+ ATPase motor^30–33^. Using ATP-dependent conformational changes in the pore motifs, Pex1–Pex6 extracts Pex5 from the peroxisome back to the cytosol, during which the TPR domain of Pex5 is thought to unfold, facilitating the release of its cargo in the peroxisomal matrix and thereby driving directionality of the import cycle^29^. In the case of the PTS2 receptors Pex18/21, extraction would dissociate both cargo and Pex7.

Currently, the mechanism by which the Pex2-10-12 complex recognizes and translocates import receptors remains largely unknown. In addition, it is unclear whether Pex2-10-12 employs a gating mechanism or forms a constitutively open pore as previously proposed^23^. The latter question is particularly relevant to the permeability properties of the peroxisomal membrane. While several lines of evidence generally support non-selective permeation of small (<∼500 Da) molecules across the peroxisomal membrane^34,35^, the presence of metabolite transporters (e.g., PMP34/Ant1) suggest certain selective properties of the permeability barrier^36,37^. Whether and how the Pex2-10-12 pore contributes to these permeability properties remains an open question.

To address these questions, here we determined cryo-EM structures of the Pex2-10-12 complex from *Saccharomyces cerevisiae* in association with Pex8, another essential factor for peroxisomal protein import in yeast. Our structural and biochemical data show that Pex8 binds to the matrix side of Pex2-10-12 and chaperones Pex5 onto Pex2-10-12 for retro-translocation. Disrupting either the Pex2-10-12–Pex8 interaction or the Pex8–Pex5 interaction impairs peroxisomal cargo import. Importantly, the structures reveal two distinct gating states of Pex2-10-12: a closed, idle conformation and an open conformation featuring a widened cleft that allows the N-terminus of Pex5 to enter. Our data indicate that the Pex5 N-terminus initially inserts into the pore as a loop. The transition from closed to open states also repositions the RING domain, enabling a ubiquitin-conjugating E2 protein to access the Pex10-RF domain only in the open conformation. Lastly, we systematically characterized sequence requirements within the Pex5 N-terminal region for pore insertion and subsequent extraction by the Pex1–Pex6 ATPase motor. Together, our study provides a detailed mechanistic framework for how the peroxisomal membrane-bound ubiquitin E3 ligase enables retro-translocation of import receptors.

## RESULTS

### Cryo-EM analysis of *S. cerevisiae* Pex2-10-12 in association with Pex8

*S. cerevisiae* has served as one of the most extensively studied model systems for peroxisomal protein import, as many Pex proteins, including Pex2-10-12, are conserved between humans and yeast. To facilitate mechanistic studies of import receptor recycling, we sought to determine the cryo-EM structure of yeast Pex2-10-12. We first purified yeast Pex2-10-12 by co-overexpression of its subunits in yeast followed by affinity purification using a SPOT-tag attached to the C-terminus of Pex12. The purified complex was eluted as a monodisperse peak in size-exclusion chromatography with the expected 1:1:1 subunit stoichiometry (Supplemental Fig. 1A).

Pex2-10-12 alone is a rather small membrane protein complex, and this limited resolution to ∼10 Å in our initial cryo-EM analysis (Supplemental Fig. 1B). The previous study of the *T. thermophilus* complex (*Tt*Pex2-10-12) used an antibody fragment to overcome this hurdle^23^, but no such tool is available for yeast Pex2-10-12. To enable high-resolution analysis, we instead searched for a native binder. Pex8 emerged as a promising candidate because Pex2-10-12 was reported as one of the interactors of Pex8 (ref. ^38^). AlphaFold modeling of Pex2-10-12 and Pex8 hinted at a potential stable interaction between them (Supplemental Fig. 1C).

Pex8 is a 65-kDa soluble peroxisomal matrix protein of undefined function. Although Pex8 is not generally found outside fungal species, it is essential for peroxisomal import in yeast^39^. We could purify yeast Pex8 from *E. coli* as a fusion to the maltose-binding protein (MBP) (Supplemental Fig. 1D). In an in-vitro binding assay, both MBP-Pex8 and cleaved Pex8 stoichiometrically associated with Pex2-10-12 (Fig. 1A). Cryo-EM analysis of Pex2-10-12 bound to MBP-Pex8 yielded two structures at overall resolutions of 2.9 and 3.1 Å (Supplemental Fig. 1E–K and Supplementary Table 1). The two structures are different in the conformations around the putative entrance for receptor retro-translocation and the RING domain (see below). Based on the presence and absence of a pore, we will refer to them as ‘closed’ and ‘open’ states. We will first use the closed structure to discuss newly identified features of the Pex2-10-12–Pex8 complex as well as biological functions of Pex8 and then compare the closed and open states.

**Figure 1.**
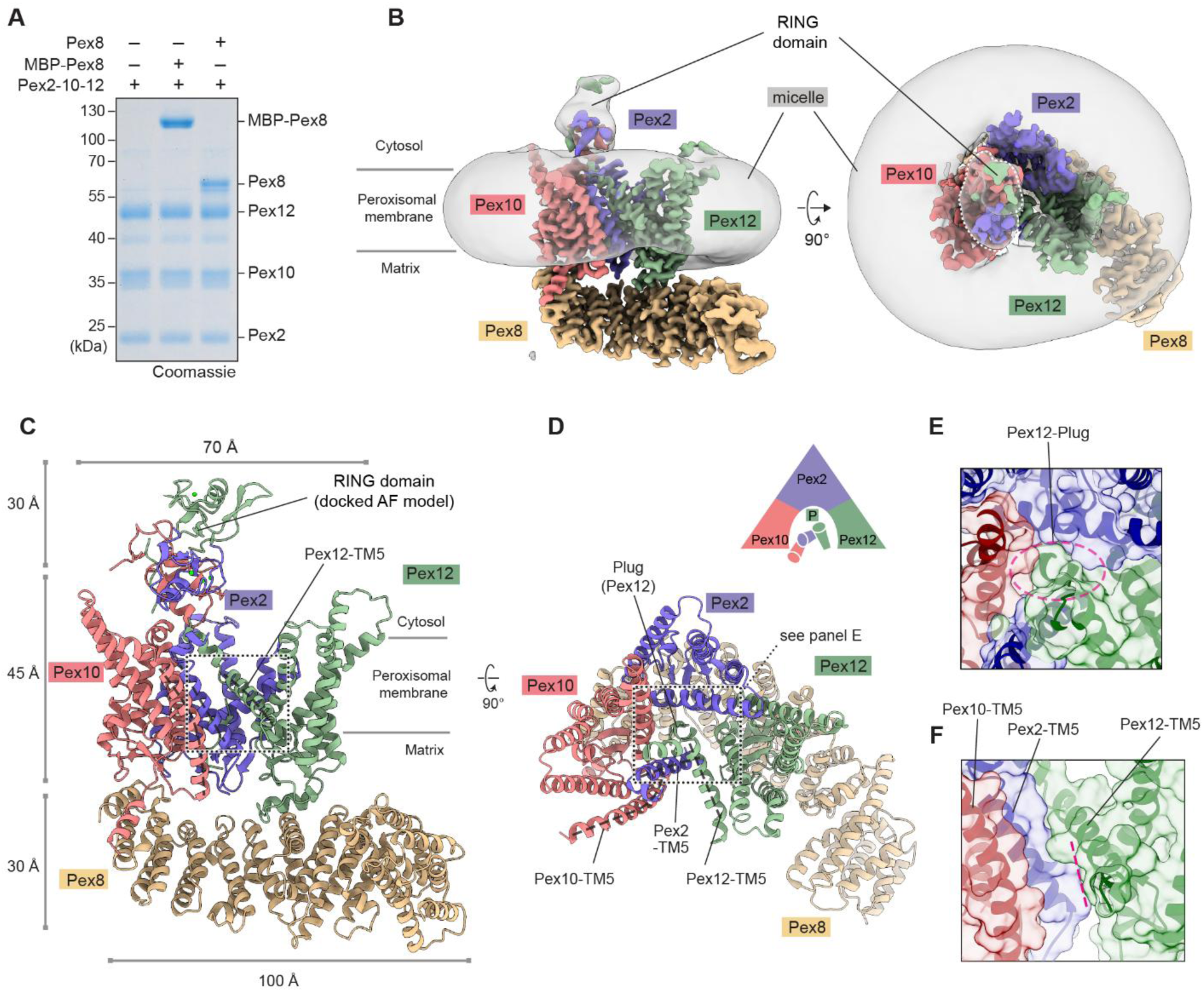
Cryo-EM structure of the yeast Pex2-10-12 complex in the closed state. (**A**) Pulldown experiment testing the interaction between Pex2-10-12 and Pex8. Agarose beads immobilized with yeast Pex2-10-12 were incubated with purified Pex8 and MBP-Pex8, washed, and the bound fractions analyzed by SDS-PAGE. (**B**) 2.9-Å-resolution cryo-EM map of Pex2-10-12 bound to MBP-Pex8 in the closed state. The high-resolution map (colored) is overlaid with a lowpass-filtered map (gray) displaying the detergent micelle and the RING domain. Left, side view; right, top view from the cytosol. (**C**, **D**) As in B, but showing the atomic model. Dashed lines indicate TM5 helices. Close-up views of the boxed regions are shown in E and F. In D, the schematic diagram illustrates the subunit arrangement with cylinders representing TM5 helices. P, plug. (**E**) Top view of the plug region. The area outlined by the magenta dashed oval is sealed by the plug. (**F**) Side view of the lateral seam region. The seam, indicated by magenta dashed line, opens in the open state (see Fig. 4D).

### Closed-state structure of the Pex2-10-12 ligase in complex with Pex8

Pex2, Pex10, and Pex12 are distantly related proteins, and each contains five transmembrane segments (TMs), the first four of which (TMs 1–4) adopt the same fold arranged into an overall shape of a baseball glove (Supplemental Fig. 2A). Pex8 is a HEAT repeat protein, and its repeated short ɑ-helices form an arch-shaped ɑ-solenoid. Our cryo-EM structure shows a well-resolved TMD of the heterotrimeric Pex2-10-12 complex and almost the entirety of Pex8 (Fig. 1B; Supplemental Fig. 2B). The C-terminal half of Pex8 is docked onto Pex2-10-12 on the matrix side with the concave side of the ɑ-solenoid facing towards the vertical axis of Pex2-10-12 (Fig. 1B–D). The RING domain is, however, visible as a low-resolution feature, likely due to some positional flexibility of the RING domain. Although the resolution of this feature was insufficient for de-novo atomic model building, its AlphaFold model could be readily fitted into the density map, allowing us to confidently determine the position and orientation of the domain (Supplementary Fig. 2B).

Viewed from the cytosol, the TMDs of Pex2-10-12 are arranged in a triangular shape with TMs 1–4 of Pex10, Pex2, and Pex12 positioned at each corner in clockwise order (Fig. 1D). The concave face of each subunit faces inward, together forming a larger cytosolic funnel. Three TM5 helices, one each from Pex2, Pex10, and Pex12, are clustered on the bottom side of the triangle. This overall TMD organization is similar to that of *Tt*Pex2-10-12 (ref. ^23^). However, structural alignment revealed a considerable deviation in intersubunit positioning: when Pex12 was superimposed, the Pex10 subunit was offset by an ∼10° rotation (Supplementary Fig. 2C). In addition, comparisons of individual subunits between the two structures showed substantial conformational differences in the TM5 helices, whereas TMs 1–4 were largely superimposable (Supplementary Fig. 2A). These observations suggest structural flexibility in the interface between Pex10 and Pex12, presumably due to loose packing among TM5 helices, which together form the interface.

One interesting feature of Pex2-10-12 is a highly tilted TM5 of Pex12, which intercalates between TM5 of Pex2 and TM1 of Pex12 (Fig. 1C,D). Almost the entire part of Pex12-TM5 forms an amphipathic helix, which bends the matrix leaflet of the membrane near the protein-lipid interface, thinning the lipid bilayer therein (Supplementary Fig. 2D). As discussed below, this region forms a putative retro-translocation pore.

Contrary to the previous *Tt*Pex2-10-12 structure, yeast Pex2-10-12 does not exhibit a constitutively open pore (Fig 1C–F). Our closed structure shows two conduit features (vertical path and lateral seam) for polypeptide translocation in the TMD, but both are occluded (Fig. 1E,F). The vertical path is blocked by a ‘plug’-like feature formed by the N-terminal segment (amino acids 1-13) of Pex12 (Fig. 1E; we note that the *Tt*Pex2-10-12 study annotated an unrelated region of Pex10 as a plug, but this feature does not gate the pore). The lateral seam, which is formed immediately below the amphipathic Pex12-TM5, is also obstructed by tight packing between Pex2-TM5, Pex12-TM1, Pex12-TM5, and a small ɑ-helix following Pex12-TM2 via highly conserved contacts (Supplementary Fig. 2E–G). In the open structure of yeast Pex2-10-12, the lateral seam widens and merges with an opening that forms in part of the vertical path to create the retro-translocation pore (see below).

### Interaction between Pex2-10-12 and Pex8 is required for peroxisomal import in yeast

The position of Pex8 in the complex suggests that it may function as a docking factor for peroxisomal import receptors during the recycling process, and without Pex8, receptors may fail to engage Pex2-10-12 efficiently. To test this hypothesis, we disrupted the interaction between Pex8 and Pex2-10-12 and examined its effects on peroxisomal protein import. Pex8 interacts with Pex2-10-12 at two sites, designated Interfaces 1 and 2, formed on the matrix loops of Pex10 and Pex12, respectively (Fig. 2A). Interface 1 is stabilized primarily by hydrophobic interactions (F20, L103, and I187 from Pex10; and W562, I574, and Y578 from Pex8), whereas Interface 2 consists of both hydrophobic (L115, L117, and L121 from Pex12; and Y340, I350, and V351 from Pex8) and polar contacts (N114 and T118 from Pex12; and D343 and T347 from Pex8).

**Figure 2.**
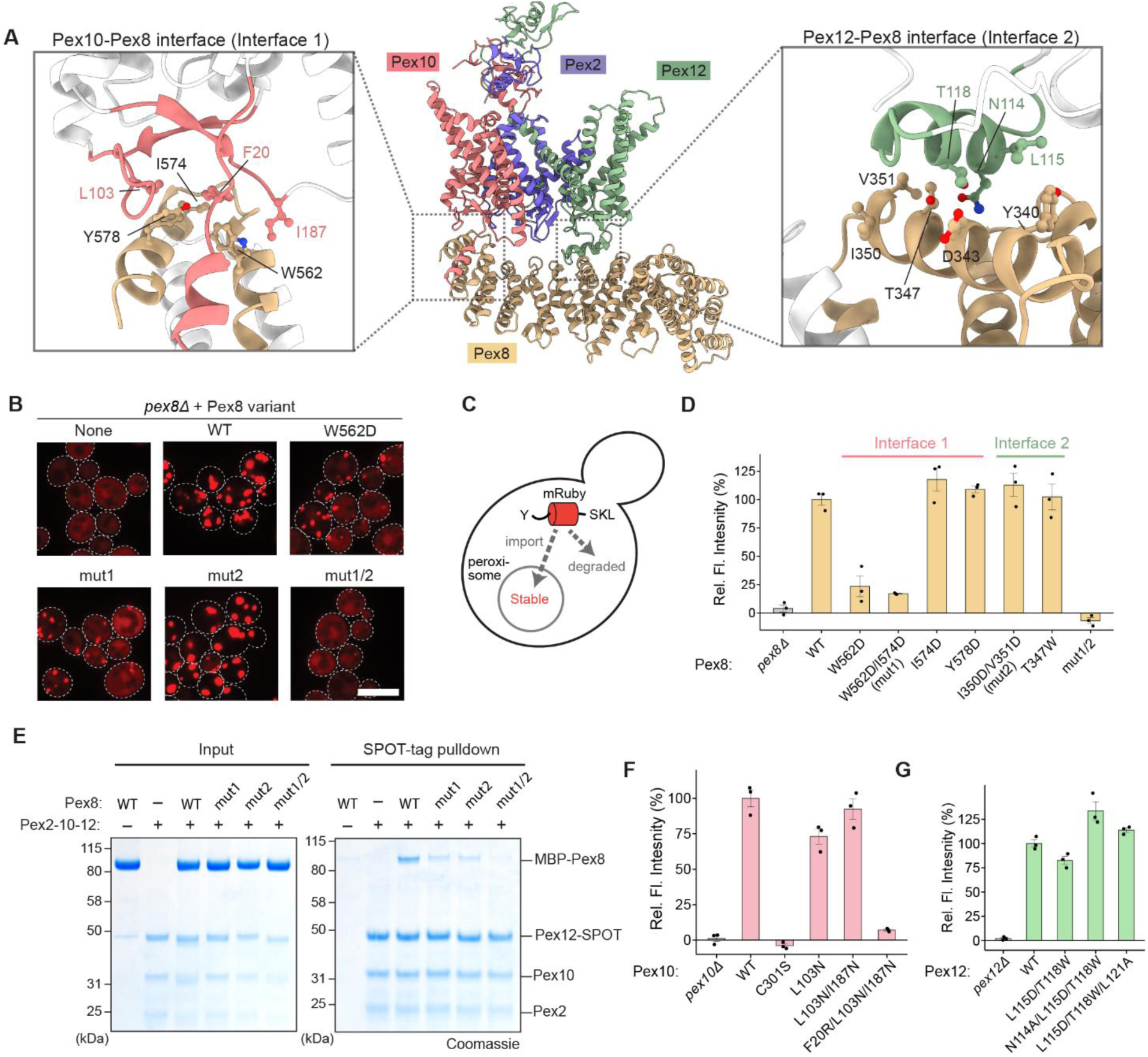
Interaction between Pex2-10-12 and Pex8 is required for peroxisomal protein import in yeast. (**A**) Close-up views of the Pex10–Pex8 and Pex12–Pex8 interfaces observed in the cryo-EM structure. Amino acid side chains at the contact sites are shown in ball-and-stick representation. (**B**) Confocal fluorescence microscopy assay testing peroxisomal import of mRuby-PTS1. Indicated Pex8 variants were expressed in *pex8Δ* yeast. ‘mut1’, W562D/I574D; ‘mut2’, I350D/V351D; ‘mut1/2’, I350D/V351D/W562D/I574D. Scale bar, 5 μm. (**C**) Schematic diagram of the fluorescence-based Ub-Y-mRuby-PTS1 reporter assay^40^. Y, N-terminally exposed tyrosine; SKL, PTS1 sequence. (**D**) Peroxisomal import efficiency of Pex8 mutants measured by the Ub-Y-mRuby-PTS1 reporter. Indicated mutants were expressed in *pex8Δ* yeast. Relative fluorescence intensities were normalized to the blank (0%) and the mean value from WT Pex8 (100%). Data represent means ± s.e.m. from three independent experiments. (**E**) In-vitro pull-down assay as in Fig. 1A but testing Interface 1 and Interface 2 mutants of Pex8. (**F**, **G**) As in D, but testing Pex10 Interface-1 mutants expressed in *pex10Δ* yeast (F) or Pex12 Interface-2 mutants expressed in *pex12Δ* yeast (G). Pex10 C301S disrupts its RF domain function. Data represent means ± s.e.m. from three independent experiments.

We first introduced point mutations into the Pex8 interfaces and tested their ability to rescue peroxisomal import defects in a *pex8Δ* strain. Two complementary peroxisomal import assays were employed: (1) a fluorescence microscopy with a red fluorescence protein (mRuby) fused to PTS1, which visualizes punctate peroxisomal structures upon successful import but remains cytosolic when import is defective (Fig. 2B), and (2) a fluorescence intensity measurement assay with a ‘Ub-Y-mRuby-PTS1’ reporter (Fig. 2C,D). In the latter assay, inefficient import leads to prolonged cytosolic retention of the reporter, triggering ubiquitin (Ub) cleavage and proteasomal degradation via the N-end rule pathway (‘Y’ denotes tyrosine), thereby decreasing cellular mRuby fluorescence^40^. Notably, these assays provide complementary dynamic ranges: the Ub-Y-mRuby-PTS1 assay is more sensitive to mild import defects due to competing N-end rule-dependent degradation, whereas the mRuby-PTS1 assay enables detection of residual import activity in strongly impaired mutants via punctate signals.

In both assays, Pex8 mutant ‘mut2’, which targets Interface 2, effectively rescued peroxisomal import deficiency of *pex8Δ*, comparable to wild-type (WT) Pex8 (Fig. 2B, Supplementary Fig. 3A). In contrast, Interface 1 mutants W562D and ‘mut1’ exhibited substantial import defects. The combined mutant (‘mut1/2’) exacerbated these defects, nearly abolishing import entirely. In-vitro pulldown experiments revealed that mut1 and mut2 partially reduced Pex8 binding to Pex2-10-12, whereas mut1/2 showed no significant interaction (Fig. 2E). These results indicate that both interfaces contribute to Pex8 docking, although disrupting Pex8–Pex12 alone has minimal functional consequences.

To determine whether Pex10 or Pex12 mutations in the interfaces also affect peroxisomal protein import, we performed complementation experiments using *pex10Δ* and *pex12Δ* strains (Fig. 2F,G; Supplementary Fig. 3B–D). While various Interface 2-targeting Pex12 mutants showed no major import defects, an Interface 1-targeting triple-point mutant (F20R/L103N/I187N) of Pex10 showed a severe reduction in import efficiency in both mRuby-PTS1 and Ub-Y-mRuby-PTS1 assays. Collectively, these results demonstrate that the Pex8–Pex10 interaction is particularly critical for peroxisomal import.

### Pex8 guides the N-terminus of Pex5 to Pex2-10-12

During recycling of Pex5 back to the cytosol, the N-terminus of Pex5 is expected to first engage Pex2-10-12 from the matrix side. Our structure suggests that Pex8 may facilitate this step by guiding the N-terminal segment of Pex5 into the retro-translocation pore of Pex2-10-12. Although our structure does not include Pex5, AlphaFold predicts that residues 49–73 of Pex5 forms two short consecutive ɑ-helices (ɑ2 and ɑ3; residues 49–53 and 63–73, respectively) and these interact with the middle section of Pex8’s concave surface (the two contact sites are referred to as Interfaces 3 and 4) (Fig. 3A,B). This segment lies downstream of a potential helical region (ɑ1), the region that was shown to be functionally important in metazoan Pex5 (ref. ^28^), and upstream of the first Pex13-interacting WxxxF/Y motif^14,17,29^ (ɑ5; residues 120–124) (Fig. 3B). The role of this interposed segment has so far not been established. In the AlphaFold model, ɑ2 and ɑ3 adopt amphipathic structures with their hydrophobic faces oriented toward Pex8 (Fig. 3A).

**Figure 3.**
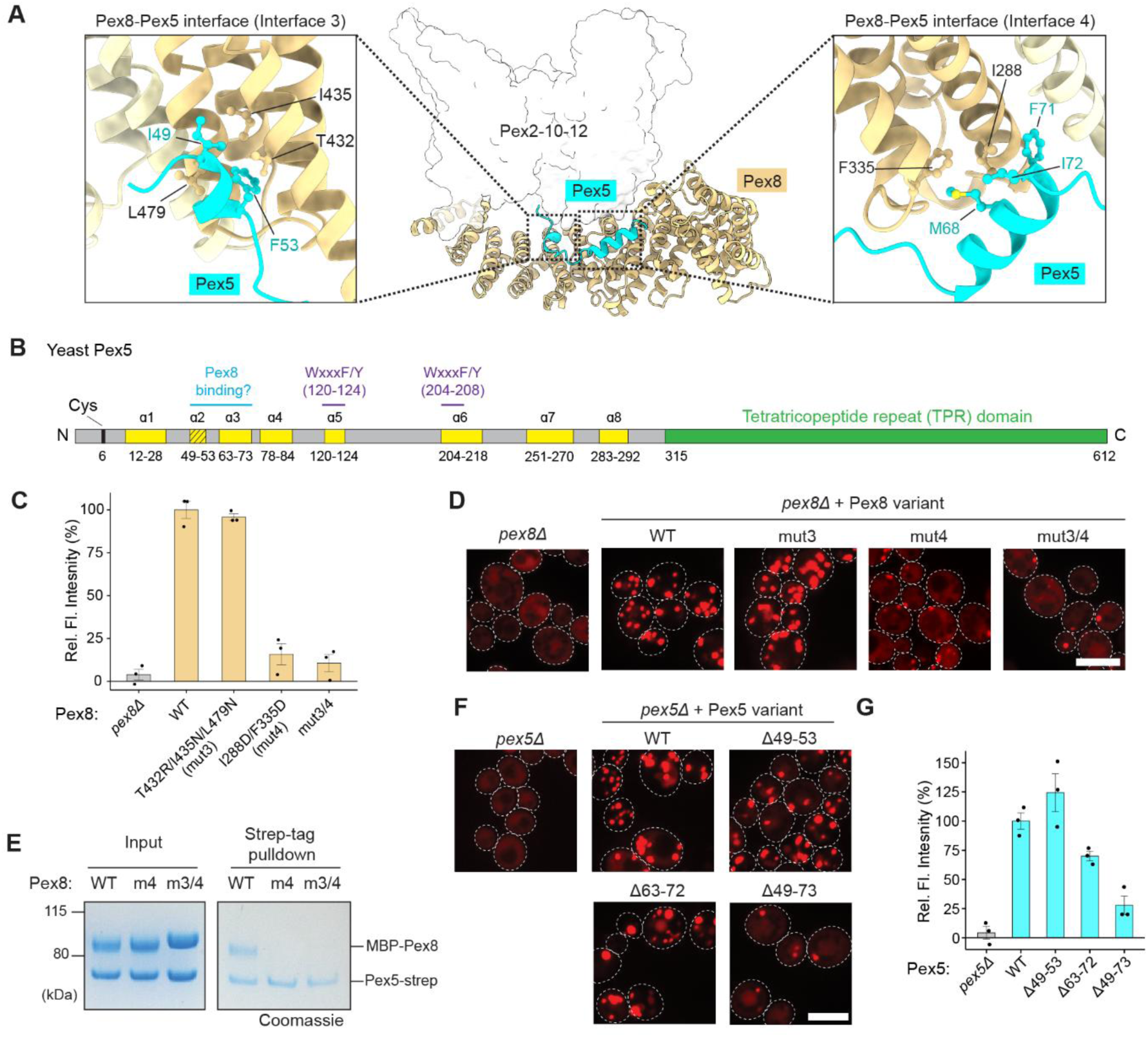
Pex8 chaperones Pex5 to the Pex2-10-12 complex. (**A**) AlphaFold-predicted interaction between the yeast Pex5 N-terminal domain and Pex8 (ref. ^58^; DOI:10.5452/ma-bak-cepc-0713). For clarity, only residues 46–74 of Pex5 are shown. Two contact sites mediated by Pex5 segments 49–53 and 63–73 are designated as Interfaces 3 and 4, respectively. The position of Pex2-10-12 is outlined based on the cryo-EM structure. (**B**) Schematic diagram of yeast Pex5. Segments highlighted in yellow form α-helices based on AlphaFold predictions. Resides 49–53 (α2) forms an α-helix only in the presence of Pex8 in the prediction. WxxF/Y motifs mediate binding to Pex13 during initial translocation across the peroxisomal membrane. (**C**, **D**) Functionality of Pex8 mutants tested by the Ub-Y-mRuby-PTS1 reporter assay (C) and mRuby-PTS1 microscopy assay (D). ‘mut3/4’: combinatorial mutant (i.e., I288D/F335D/T432R/L479N/I435N). In C, data represent means ± s.e.m. from three independent experiments. In D, scale bar, 5 μm. (**E**) In-vitro pull-down assay with purified Strep-tagged Pex5 and MBP-Pex8. ‘m4’: mut4; ‘m3/4’: mut3/4. (**F**, **G**) As in C and D, but testing Pex5 truncation mutants. Pex5 is attached to a C-terminal HA (in F) or ALFA (in G) tag. In G, data represent means ± s.e.m. from three independent experiments.

To test the functional importance of this predicted interaction between Pex8 and Pex5, we first mutated the Interface 3 and 4 regions in Pex8 and tested their effects on peroxisomal protein import (Fig. 3C,D; Supplementary Fig. E,F). In both Ub-Y-mRuby-PTS1 and microscopy-based import assays, Interface 3 mutant (mut3) showed little or mild defects, whereas an Interface 4 mutant (mut4) and the combined mutant (mut3/4) showed strong import defects. In an in-vitro pulldown assay, Pex8 with mut4 alone already barely copurified with Pex5, consistent with its poor ability to support peroxisomal import in cells (Fig. 3E). Taken together, these data demonstrate that Pex5 binding to Interface 4 of Pex8 is crucial for Pex5 recycling.

Next, we truncated the segment in Pex5 that is predicted to interact with Pex8 in Interfaces 3 and 4, and tested the ability of the mutants to rescue import defects in a *pex5Δ* strain. In the fluorescence microscopy assay, both Δɑ2 (Δ49–53) and Δɑ3 (Δ63–72) Pex5 mutants supported the peroxisomal localization of the mRuby-PTS1 reporter like WT Pex5 (Fig. 3F; Supplementary Fig. 3J). However, the Δɑ2/3 (Δ49–73) mutant removing both regions showed a noticeable reduction in peroxisomal localization of mRuby-PTS1 with a concomitant increase in cytosolic localization. In addition, the Ub-Y-mRuby-PTS1 assay showed more pronounced import defects with the Δɑ2/3 mutation (Fig. 3G). These results, together with strongly defective phenotypes of mut3/4 Pex8, support the Pex8–Pex5 interaction predicted by AlphaFold. However, we note that Δɑ2/3 Pex5 exhibits substantial residual peroxisomal localization of mRuby-PTS1 unlike *pex5Δ*, and the Δɑ3 mutant shows a considerably weaker phenotype than that of Pex8 mut4. This suggests that the interaction between Pex5 and Pex8 does not strictly depend on the pattern that AlphaFold predicts, and other Pex5 N-terminal regions may be able to partially substitute the role of ɑ2 and ɑ3 in interacting with Pex8. Additionally, the TPR domain of Pex5 may also contribute to Pex8 binding as Pex8 contains a PTS1 at its C-terminus.

### Open Pex2-10-12 structure reveals the gating mechanism

In our single-particle cryo-EM analysis, about 54% and 46% particles were classified into closed and open states, respectively (Supplementary Fig. 1F). In both structures, Pex2-10-12 is associated with Pex8 in essentially the same manner through Interfaces 1 and 2 (Fig. 4A–B). TMs 1–4 of Pex2, Pex10, and Pex12 also maintain the overall structure with some moderate movements in TMs 2–4 of Pex10 and TMs 1 and 2 of Pex12 (Fig. 4C; Supplementary Fig. 4A). By contrast, the three TM5s show substantial rearrangements, leading to several important conformational changes in the TMD, as described below.

**Figure 4.**
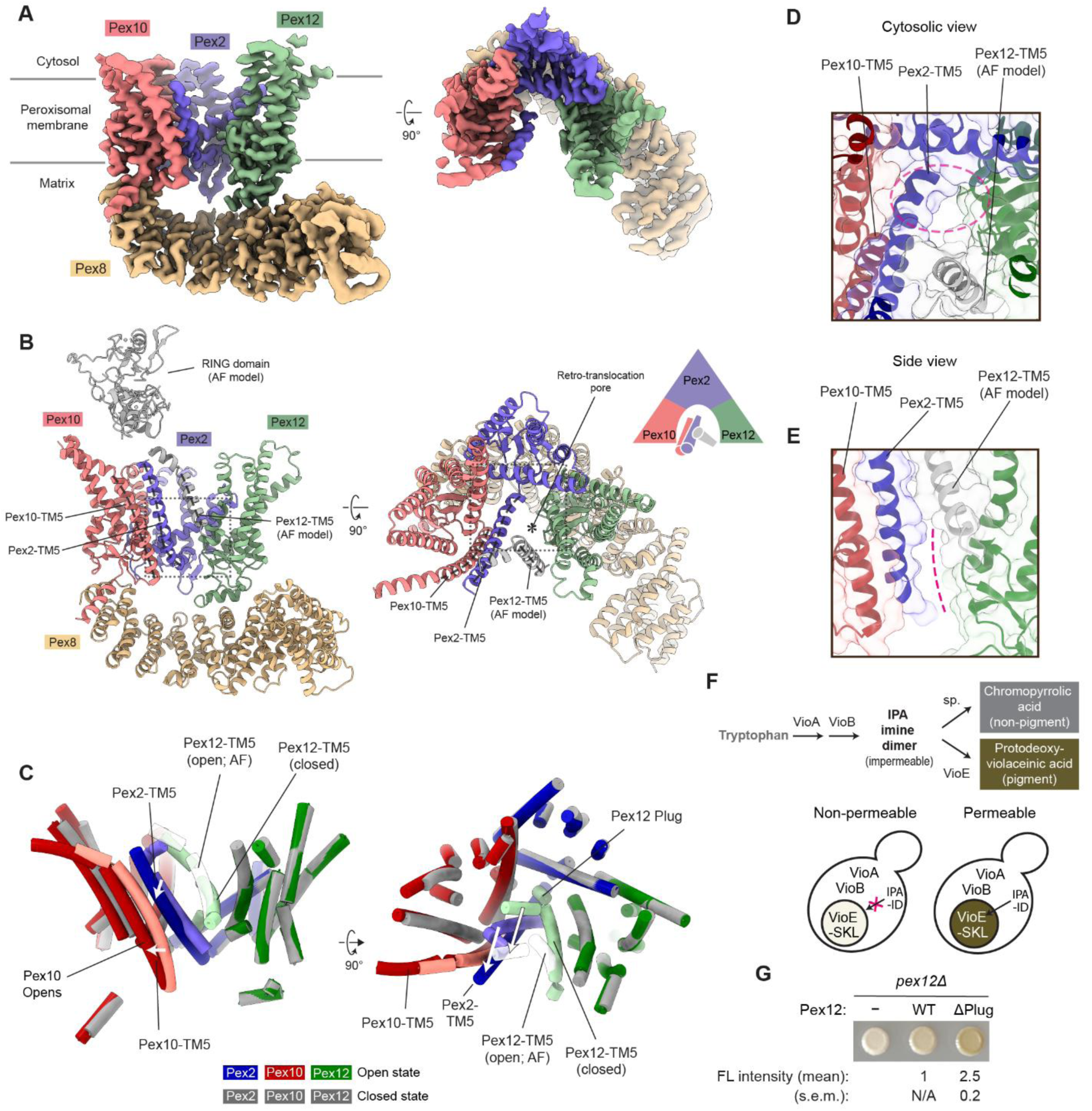
Cryo-EM structure of the yeast Pex2-10-12 complex in the open state. (**A**) Cryo-EM map of yeast Pex2-10-12–Pex8 in the open state at 3.1 Å resolution. Left, side view; right, the top view from the cytosol. (**B**) As in A, but showing the atomic model. The RING domain and TM5 of Pex12 (gray) were modeled using AlphaFold due to poor cryo-EM density. Dashed lines indicate TM5 helices. Close-up views of the boxed regions are shown in C and D. (**C**) Conformational changes in the TMD accompanied by the closed-to-open transition of Pex2-10-12. (**D**) Top view of the retro-translocation pore region. The region outlined by the magenta dashed oval is sealed by the plug in the closed state (compare with Fig. 1E). (**E**) Side view of the lateral seam. The magenta dashed line highlights the open seam (compare with Fig. 1F). (**F**, **G**) Violacein intermediate production assay adopted for testing permeation of indole-3-pyruvic acid imine dimer (IPAID) across the peroxisomal membrane. Upon entering the peroxisomal matrix, IPAID is converted by VioE into a dark green pigment. In G, WT or plug-deleted (Δ2-26; ΔPlug) Pex12 was expressed in *pex12Δ* yeast co-expressing VioA, VioB, and Ub-R-VioE-PTS1. Yeast cells were spotted, and their pigment levels were measured by fluorescence (FL). Intensity values were background-subtracted using non-Pex12-expressing (‘−’) cells and normalized to the value from WT Pex12-expressing cells. Data represent means ± s.e.m. from three independent experiments.

First, in the open structure, the plug in the vertical path is displaced and becomes invisible (Fig. 4D). Given the membrane topology of Pex12-TM1, the plug is likely pushed out toward the matrix, where it remains flexibly tethered. Notably, the plug displacement does not simply vacate the space it occupies in the closed state to create an open pore. Instead, this space becomes largely filled by the segment immediately preceding TM5 of Pex2, which extends the Pex2 TM5 helix (Fig. 4D; compare with Fig. 1E). This rearrangement is accompanied by a tilting of Pex2-TM5 and a shift in Pex10-TM5 (Fig. 4C).

Second, the tilting of Pex2-TM5 widens the lateral seam formed between Pex2-TM5 and Pex12-TM1, creating a wedge-shaped opening (Fig. 4A,E). The lower end of this opening extends into part of the space freed by plug displacement. The upper end should be constrained by Pex12-TM5 (Fig. 4B,E). However, in the cryo-EM map of the open state, Pex12-TM5 is barely visible, unlike the closed structure, suggesting increased flexibility in this region (Fig. 4A; compare with Fig. 1B). Consistent with this observation, Pex12-TM5 is one of the lowest-confidence regions in an AlphaFold model, which otherwise closely matches our cryo-EM structure of the open state (Supplementary Fig. 4B,C).

Our structural findings on the gating mechanism of the Pex2-10-12 complex have important implications for the permeability barrier. Although the peroxisomal membrane is substantially permeable to small ions and metabolites, it excludes molecules larger than ∼500–700 Da. We hypothesized that the plug might limit leaky permeation when the pore is not engaged in receptor retro-translocation. To test this, we adapted the previously developed protodeoxyviolacein (PDV) production assay^35^ and examined the entry of the membrane-impermeable intermediate indole-3-pyruvic acid imine dimer (IPAID; MW = 400 Da) into the peroxisomal matrix (Fig. 4F). The assay utilizes three bacterial enzymes, VioA, VioB, and VioE, that mediate successive PDV biosynthesis reactions. VioA and VioB localize to the cytosol, whereas VioE is targeted to the peroxisomal matrix by a C-terminal PTS1 in import-competent cells, thereby spatially separating the VioE-mediated PDV-producing reaction from its cytosolic IPAID substrate generated by VioA and VioB. To eliminate background signal, we fused VioE-PTS1 with a strong N-degron (Ub-R; ‘R’ denotes arginine) to deplete the cytosolic pool of VioE-PTS1, preventing pigment formation even when peroxisomal import is blocked (Fig. 4G, empty vector). In this configuration, PDV pigment is produced only if cytosolic IPAID diffuses into the peroxisomal matrix where VioE-PTS1 resides. We then compared pigment levels after expressing either WT Pex12 or a plug-deleted mutant (ΔPlug). Both constructs restored import activity in separate fluorescence-based assays confirming that the plug is dispensable for retro-translocation (Supplementary Fig. 4D,E). However, ΔPlug Pex12 yielded 2.5-fold higher pigment levels than WT (Fig. 4G), indicating that the plug removal increases IPAID permeation through the open pore. These results support the model that gating reduces leakage of certain small molecules across the peroxisomal membrane.

### Conformational switching of the RING domain during the closed-to-open transition

Another major conformational difference between the closed and open states involves the position of the RING domain (Fig. 5A,B). In the open structure, the RING domain density is much weaker and poorly resolved compared to the closed structure (Supplementary Fig. 5A), suggesting increased flexibility. Nevertheless, its position aligns remarkably well with the position predicted by AlphaFold (Supplementary Fig. 5B), thus providing a plausible model for the conformational shift. In the closed state, the RING domain is positioned near the vertical axis of the Pex2-10-12 TMD (Fig. 1B,C). Upon transitioning to the open state, the domain undergoes a rigid-body movement of ∼30–40 Å towards the Pex10 corner (Fig. 5A,B). This movement is likely driven by a conformational change in the Pex12-TM5, which connects to the RING domain through a stem-like helical segment (amino acid positions 296 to 306) (Supplementary Fig. 5C). As described below, an important consequence of this repositioning is the altered accessibility of Pex10-RF for binding to a ubiquitin-conjugated E2 (E2∼Ub).

**Figure 5.**
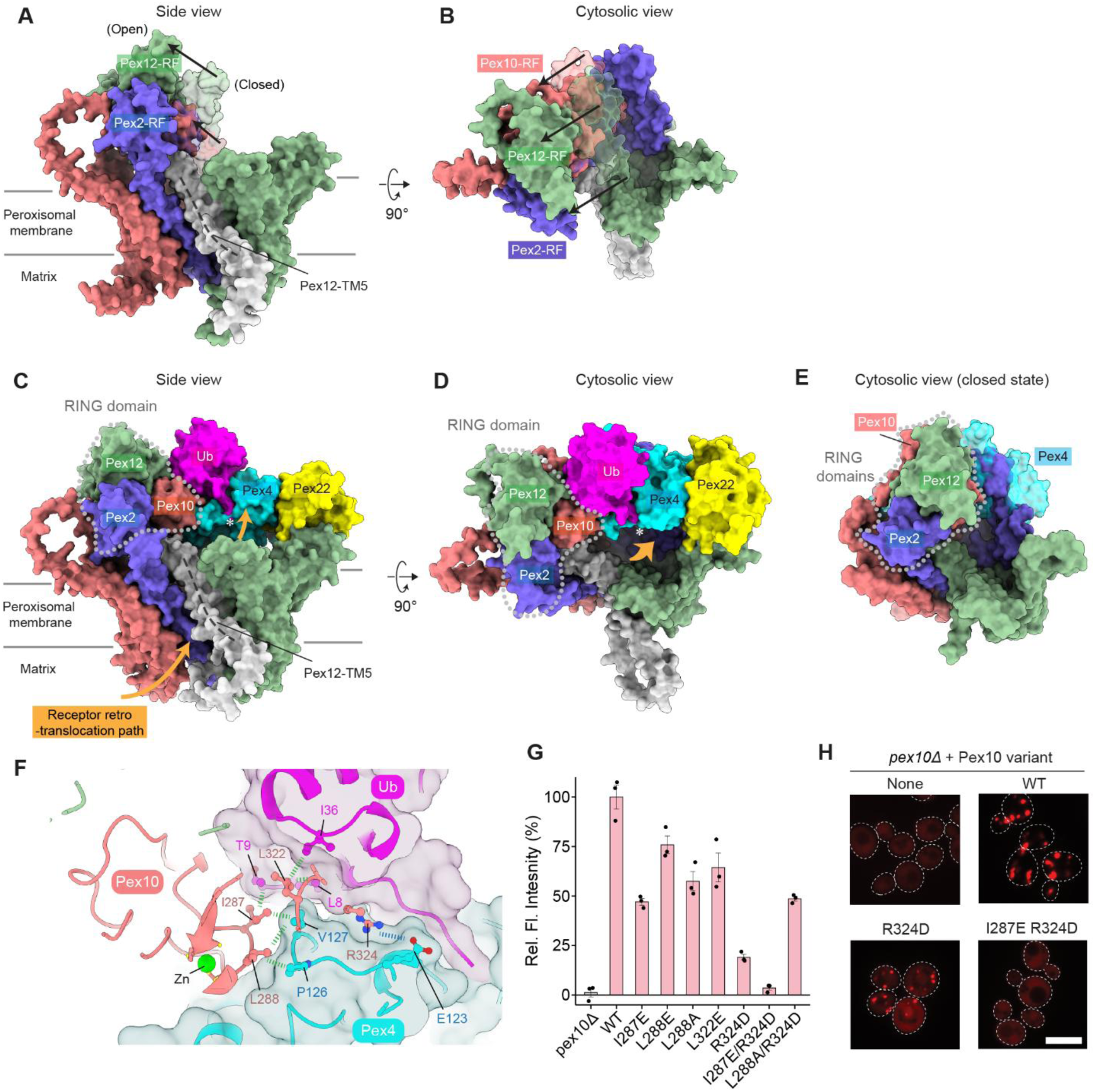
Closed-to-open transition of Pex2-10-12 regulates Pex10 Ring Finger (RF) domain for receptor ubiquitination. (**A**, **B**) Movement of the RING domain during the closed-to-open transition. The open-state AlphaFold model (solid colors) is aligned with the closed-state cryo-EM structure, with the closed-state RING domain shown as a semi-transparent surface. (**C**, **D**) AlphaFold prediction of Pex2-10-12 bound to Pex4 (E2 ubiquitin-conjugating enzyme), Pex22, and ubiquitin (Ub). The RING domain is outlined with a gray dotted line. Orange arrows indicate putative retro-translocation path. Asterisk, Ub C-terminus. (**E**) Similar to D, but showing the structure in the closed state. Pex4 (semi-transparent cyan), docked onto Pex10-RF according to the AlphaFold prediction, exhibits major clashes with the Pex2 TMD. For clarity, Pex22 and Ub are not shown. (**F**) Predicted interface among Pex10-RF, Pex4, and ubiquitin. Green dashed lines indicate putative interactions between amino acids. (**G**) Ub-Y-mRuby-PTS1 reporter assay testing peroxisomal import defects caused by Pex10-RF mutations. Data represent means ± s.e.m. from three independent experiments. (**H**) As in G, but fluorescence microscopy assay using the mRuby-PTS1 reporter. Scale bar, 5 μm. Also see Supplementary Fig. 6G.

Typically, RF domains interact with an E2∼Ub through their two Zn^2+^-coordinating loops^41–43^. As noted previously^23^, although Pex2-10-12 has three RF domains, only Pex10-RF possesses canonical features for E2∼Ub binding. Pex12-RF lacks one of the two Zn^2+^-binding motifs, and Pex2-RF is missing key residues for the E2∼Ub interaction. In addition, in the Pex2-RF, the loop linking it to TM5 sterically blocks the putative E2∼Ub binding surface according to the AlphaFold model. Consistent with these observations, AlphaFold predicts high-confidence RF–E2∼Ub assemblies only for Pex10-RF, with the primary E2 enzyme Pex4 or auxiliary enzyme Ubc4 (ref. ^25,44,45^) (Supplementary Fig. 5D–P).

In yeast and plants, Pex4 is anchored to the peroxisomal membrane by Pex22, which also allosterically activates Pex4 (ref. ^46–48^). To gain mechanistic insights into the coordination among these factors, we modeled the full Pex2-10-12 complex with Pex4, Pex22, and ubiquitin using AlphaFold (Fig. 5C,D; Supplementary Fig. 6A–C). In this model, Pex2-10-12 adopts a conformation consistent with the open state for both TMD and RING domain. Pex22–Pex4∼Ub docks onto the top of the TMD with Pex4∼Ub interfacing directly with the canonical E2-binding surface of Pex10-RF, whereas Pex22 positions near the Pex12 corner. Notably, the C-terminal double glycine motif of the ubiquitin is positioned immediately above the funnel of the TMD and thus optimally placed for the transfer of ubiquitin to the import receptor emerging from the retro-translocation pore. By contrast, manual docking of Pex4 onto Pex10-RF in the closed structure resulted in severe steric clashes with the TMDs of Pex2 and Pex10 (Fig. 5E; Supplementary Fig. 6D,E). These observations suggest that the RING domain is switched off in the closed state and that its activation is conformationally coupled to opening of the retro-translocation pathway.

While our structural model suggests that Pex10-RF mediates engagement with Pex4∼Ub, previous studies have reported conflicting views on which RF domain catalyzes receptor mono-ubiquitination. Some studies implicated Pex10-RF (ref. ^21,49^), whereas others proposed Pex2-RF or Pex12-RF as the catalytic site^20,23^. To test the direct role of Pex10-RF in receptor mono-ubiquitination, we introduced mutations at its predicted interface with Pex4∼Ub. The AlphaFold model predicted that Pex10 residues I287, L288, and L322 make hydrophobic contacts with Pex4 or ubiquitin, while R324 (also known as the linchpin residue) forms a salt bridge with E123 of Pex4 (Fig. 5F). Mutations at these Pex10 positions indeed caused varying degrees of import defects despite their WT-level expression (Fig. 5G,H; Supplementary Fig. 6F,G). Notably, the double mutant I287E/R324D completely abrogated peroxisomal import. These results provide strong evidence that Pex10-RF catalyzes receptor mono-ubiquitination required for retro-translocation.

### Pex5 N-terminal segment inserts into the Pex2-10-12 pore as a loop

Next, we sought to understand how the largely unstructured N-terminal domain of the Pex5 receptor initiates retro-translocation through the relatively narrow Pex2-10-12 pore. In principle, two scenarios are possible: (1) insertion initiated directly from the Pex5 N-terminus, or (2) insertion initiated as a loop, then followed by the N-terminus slipping into the cytosolic funnel of Pex2-10-12 (Fig. 6A). At present, it remains unclear which mechanism underlies Pex2-10-12-mediated retro-translocation.

**Figure 6.**
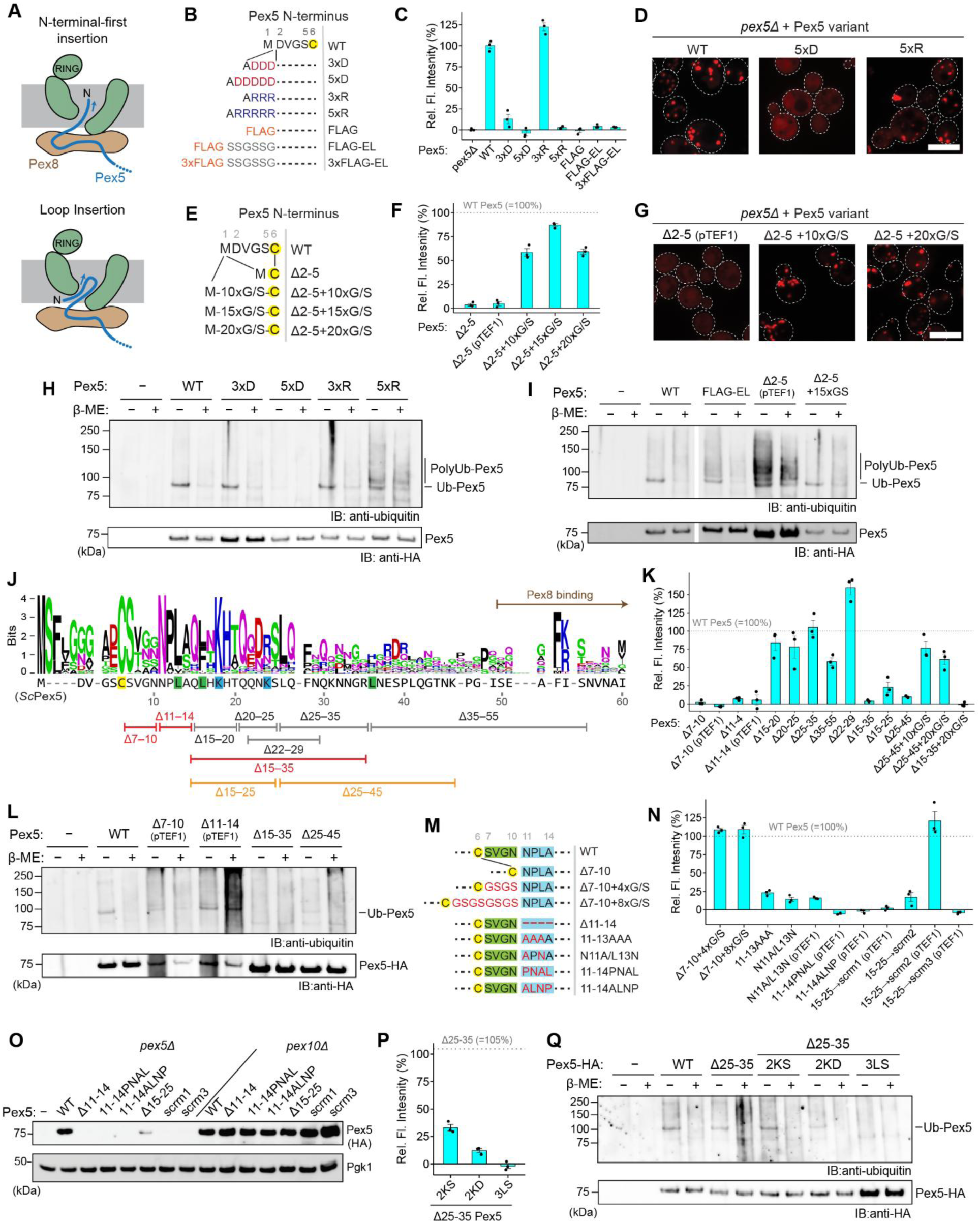
Mechanism of Pex5 insertion to the Pex2-10-12 pore. (**A**) Schematic illustration of two possible models for how the Pex5 N-terminus inserts into the Pex2–10–12 pore. (**B**) Design of Pex5 variants containing short, charged sequences appended to the N-terminus. (**C**) Ub-Y-mRuby-PTS1 reporter assay testing import activity of Pex5 variants shown in B in a *pex5Δ* background. Data represent means ± s.e.m. from three independent experiments. (**D**) Fluorescence microscopy analysis of the same Pex5 variants using mRuby-PTS1 as an import reporter. Also see Supplementary Fig. 7A. Scale bar, 5 µm. (**E**) Δ2–5 Pex5 variants carrying a 10–20 residue Gly/Ser extension at the N-terminus. (**F**, **G**) Ub-Y-mRuby-PTS1 reporter assay and fluorescence microscopy (as in C and D) testing Δ2–5 Pex5 variants with or without the Gly/Ser tail. ‘pTEF1’ indicates expression driven by the stronger *TEF1* promoter instead of the default, weaker *ALD6* promoter. In F, data represent means ± s.e.m (n=3). In (G), scale bar is 5 µm. (**H**, **I**) Ubiquitin blotting assay. C-terminally HA-tagged Pex5 variants with the indicated fusions or deletions were immunoprecipitated and probed with anti-ubiquitin antibody. Samples were treated with 2-mercaptoethanol (β-ME), where indicated. (**J**) Logo plot representing N-terminal sequence conservation among 90 fungal Pex5 clustered sequences. Cys6 and Lys18/Lys24 are highlighted in yellow and blue, respectively. Leu13, Leu16, and Leu36 are highlighted in green; mutation of these residues was tested for pore insertion (see panels O and P). Bars indicate residue ranges used for truncation variants, with colors denoting import activity: gray, functional; orange, partially defective; red, inactive. (**K**) Ub-Y-mRuby-PTS1 assay testing internal truncation variants of Pex5. Data represent means ± s.e.m (n=3). (**L**) Ubiquitin blotting assay examining ubiquitination of Pex5 truncation variants. (**M**) Pex5 variants carrying deletions, insertions, or substitution mutations within residues 7–14. (**N**) Ub-Y-mRuby-PTS1 assay testing Pex5 variants shown in (M) and sequence-scrambled versions of residues 15–25 (WT=QLHKHTQQNKS; scrm1=KKSQNTLQHHQ; scrm2= TQLNSQHKHQK; scrm3=NLKSHKQQTHQ). Data represent means ± s.e.m (n=3). (**O**) The indicated Pex5 variants were expressed in *pex5Δ* or *pex10Δ* yeast, and their steady-state levels were analyzed by immunoblotting. Pgk1, loading control. (**P**) Ub-Y-mRuby-PTS1 assay testing Δ25–35 Pex5 variants carrying the indicated additional mutations. 2KS = K18S/K24S; 2KD = K18D/K24D; 3LS = L13S/L16S/L36S. Data represent means ± s.e.m (n=3). (**Q**) Ubiquitin blotting assay testing Pex5 variants shown in (O).

To distinguish between these two possibilities, we first tested the effect of appending short amino acid sequences to the Pex5 N-terminus (Fig. 6B). We reasoned that if pore insertion begins at the N-terminus, the presence of additional segments would strongly impair retro-translocation efficiency and, consequently, the overall protein import activity of Pex5. We first examined fusions of three or five consecutive charged amino acids (3xD, 5xD, 3xR, and 5xR), as well as FLAG-tags of different lengths, which also carry multiple charged amino acids. These sequences were chosen because charged amino acids often pose energetic barriers to protein translocation.

In both Ub-Y-mRuby-PTS1 and microscopy-based assays (Fig. 6C,D; Supplementary Fig. 7A,B), we observed pronounced peroxisomal import defects for several of these Pex5 variants. For example, addition of 5xD completely abolished import, while FLAG-tagged variants also exhibited little to no import activity. By contrast, Pex5 fused to 5xR retained substantial import activity in the microscopy assay, although the Ub-Y-mRuby-PTS1 assay reported virtually no activity, suggesting that 5xR Pex5 can still undergo retro-translocation, albeit inefficiently. Furthermore, 3xR Pex5 showed no detectable import defects in either assay, indicating tolerance to a few consecutive charged residues at the N-terminus.

The PTS1 reporter-based assays measure only overall retro-translocation efficiency and therefore provide limited insight into the specific mechanistic steps underlying defective phenotypes. In addition to pore insertion, impaired engagement with the Pex1–Pex6 ATPase motor could also compromise retro-translocation. Because mono-ubiquitination occurs at Cys6, the preceding N-terminal residues (positions 1–5) are likely essential for the Pex1–Pex6 engagement, analogous to the unstructured initiation tail needed for the proteasomal processing of a polyubiquitinated substrate^50^. Fusion of highly charged sequences may have disrupted engagement with Pex1–Pex6 rather than the initial insertion into the Pex2-10-12 pore. Thus, as an alternative approach, we deleted residues 2–5 of Pex5 (Δ2–5) and appended neutral Gly/Ser stretches of varying lengths (Fig. 6E). As expected, Δ2–5 Pex5 failed to support peroxisomal import, consistent with the loss of an initiation tail for the Pex1–Pex6 motor, (Fig. 6F,G; Supplementary Fig. 7C) Although Δ2–5 Pex5 was expressed at a substantially lower level, likely due to accelerated degradation, enhanced expression with the strong *TEF1* promoter did not rescue import. Strikingly, however, adding a 10–20-residue Gly/Ser stretch to the N-terminus of Δ2–5 Pex5 largely restored import activity. This indicates that the Gly/Ser stretches block neither the pore insertion nor the Pex1–Pex6-mediated extraction. These findings thus support a loop-insertion model.

We then asked whether the inability of 5xD- or FLAG-tag-fused Pex5 to support peroxisomal import stems from defects in the initial insertion into the Pex2-10-12 pore or from impaired downstream engagement with the Pex1–Pex6 motor. To distinguish between these possibilities, we immunoprecipitated Pex5 variants and probed for their ubiquitination. Failure to insert into Pex2-10-12 would be expected to block formation of the characteristic mono-ubiquitinated species. Consistent with prior reports^25^, WT Pex5 showed a defined ubiquitinated band that disappeared after β-mercaptoethanol (β-ME) treatment, reflecting cleavage of the thioester linkage between Cys6 of Pex5 and the C-terminus of ubiquitin (Fig. 6H). In contrast, Δ2–5 Pex5 displayed a polyubiquitination pattern resistant to β-ME (Fig. 6I), consistent with the receptor accumulation and degradation in the absence of recycling (RADAR), where lysine residues (Lys18 and Lys24) of the Pex5 N-terminus are polyubiquitinated by Pex2-10-12 upon failure of extraction by Pex1–Pex6 (ref. ^51,52^). Notably, adding a 15xGly/Ser stretch to Δ2–5 Pex5 restored import activity along with a WT-like ubiquitination profile (Fig. 6I). Interestingly, although both 5xD- and FLAG-tagged Pex5 were similarly defective for import, they showed distinct ubiquitination patterns: 5xD Pex5 exhibited a dramatic reduction in ubiquitination, consistent with impaired pore insertion, whereas FLAG-tagged Pex5 accumulated polyubiquitination, indicating successful insertion but inefficient extraction (Fig. 6H). These findings suggest that while the Pex5 N-terminal region inserts into the Pex2-10-12 pore as a loop, fusion of a highly negatively charged sequence, such as 5xD, can hinder completion of the insertion (note that the FLAG-tag carries a net charge of −3, compared to −5 for 5xD).

### Defining motifs in the Pex5 N-terminal segment required for recycling

We next sought to define the roles of the Pex5 N-terminal region in retro-translocation. To this end, we systematically truncated internal segments within the first ∼50 residues of Pex5, downstream of Cys6 and upstream of the Pex8-binding region (Fig. 6J; Supplementary Fig. 7C-F). Sequence alignment of fungal Pex5 revealed moderate conservation up to residue ∼27, with a sharp decline thereafter. Most deletion variants removing subregions between residues 15–55 (Δ15–20, Δ20–25, Δ25–35, Δ35–55, and Δ22-29) did not show major import defects in the mRuby-PTS1 and Ub-Y-mRuby-PTS1 assays, indicating that short (5-residue) segments within this region are individually dispensable (Fig. 6K; Supplementary Fig. 7D, F, and G).

By contrast, four-amino-acid deletions within residues 7−14 (Δ7-10 and Δ11-14) caused severe import defects accompanied by markedly reduced steady-state protein levels (Fig. 6K; Supplementary Fig. 7D,G). Overexpression from the *TEF1* promoter did not rescue import-deficient phenotypes. Anti-ubiquitin immunoblotting further showed that both mutants are ubiquitinated at non-Cys6 positions, suggesting that their defects most likely result from inefficient Cys6 ubiquitination and subsequent extraction failure leading to RADAR-mediated degradation, rather than impaired pore insertion (Fig. 6L). Strikingly, adding back a 4- or 8-residue Gly/Ser stretch to Δ7–10 fully restored import activity, indicating that the original defect of Δ7–10 likely arose from shortening of the distance between Cys6 and a downstream sequence (Fig. 6M,N; Supplementary Fig. 7C,D). Likewise, substitution of residues 11–13 with 3xAla (11–13AAA) or the N11A/L13N mutation caused noticeably milder import defects compared to Δ11-14. However, scrambled (11–14PNAL and 11–14ALNP) mutants showed severe import impairment accompanied with markedly reduced protein levels (Fig. 6M,N; Supplementary Fig. 7D, G, and H), consistent with RADAR-mediated degradation, as their expression was restored to WT levels in the *pex10Δ* background (Fig. 6O). Collectively these data show that although not strictly sequence-dependent, residues 11–14 are required for efficient receptor mono-ubiquitination while remaining dispensable for pore insertion.

Truncation analyses suggest only a weak sequence-specific requirement in the succeeding region. However, the data also point to potential redundancy among functionally important elements. For example, while small deletions within residues 15–35 (Δ15–20, Δ20–25, and Δ25–35) were functional, a larger deletion (Δ15–35) was completely inactive (Fig. 6K; Supplementary Fig. 7E,F). Similarly, Δ25–45 was barely active. Notably, adding back a 10x or 20x Gly/Ser segment to Δ25–45 rescued import, whereas a 20xGly/Ser insertion into Δ15–35 did not. These results indicate that residues 15–35 harbor an element critical for retro-translocation. Consistent with this, no apparent ubiquitination was observed with the Δ15–35 mutation (Fig. 6L).

Unlike the inactive Δ15–35 mutant, Δ15–25 exhibited partially impaired import activity along with a reduced protein level (Fig. 6O; Supplementary Fig. 7E,H). This instability was also observed in variants where residues 15–25 were scrambled: while one mutant (scrm2) was functional, others (scrm1 and scrm3) showed only slight import activity and markedly low expression. Again, the reduced protein levels in these defective mutants are consistent with their degradation by the RADAR pathway, as their protein levels were comparable to WT in *pex10Δ*. The fact that these variants are targeted by RADAR implies that like the 11–14PNAL or ALNP mutants, they can still successfully insert into the Pex2-10-12 pore but are inefficiently mono-ubiquitinated at Cys6, potentially due to improper positioning of Cys6 with respect to the RING domain.

To identify the specific feature within residues 15–35 required for pore insertion, we leveraged the fully functional Δ25–35 Pex5 variant and introduced substitution mutations around positions 15–25 within this truncation background (Fig. 6P,Q; Supplementary Fig. 7I,J). Replacement of positively charged residues Lys18 and Lys24, which also serve as poly-ubiquitination sites in RADAR^51,52^, with serine or aspartic acid caused only partial import defects. By contrast, simultaneous substitution of three hydrophobic amino acids (L13, L16, and L36) with serine in Δ25–35 Pex5 completely abolished import activity while retaining WT-level expression. Similar to the Δ15–35 mutant, this mutant also showed no detectable Cys6 ubiquitination. Taken together, these results suggest that efficient pore insertion requires hydrophobic amino acids in this region but does not depend on a precise amino acid sequence.

### Model for the Pex2-10-12 ligase-mediated receptor recycling

Our study refines the mechanistic model of import receptor recycling mediated by the Pex2-10-12 complex (Fig. 7). After delivering cargo into the peroxisomal matrix via the Pex13–Pex14 import pore (importomer), Pex5 (or PTS2 receptor Pex18/21) is recruited to Pex8 via interactions with the central region of the Pex8 HEAT-repeat solenoid. In its resting state, the Pex2-10-12 ligase adopts a closed conformation: the retro-translocation conduit is occluded by the plug and lateral seam in closed states, and the cytosolic Pex10-RF is inaccessible for binding of the E2 enzyme Pex4. A concerted conformational switch opens the complex, displacing the plug, allowing insertion of the Pex5 N-terminus into the pore, and recruiting Pex4∼Ub to Pex10-RF. The Pex5 N-terminal segment inserts into the pore as a loop, likely mediated by hydrophobic contacts between the pore and Pex5 residues ∼11–35 (‘Pex2-10-12-interacting’ segment in Fig. 7).

**Figure 7.**
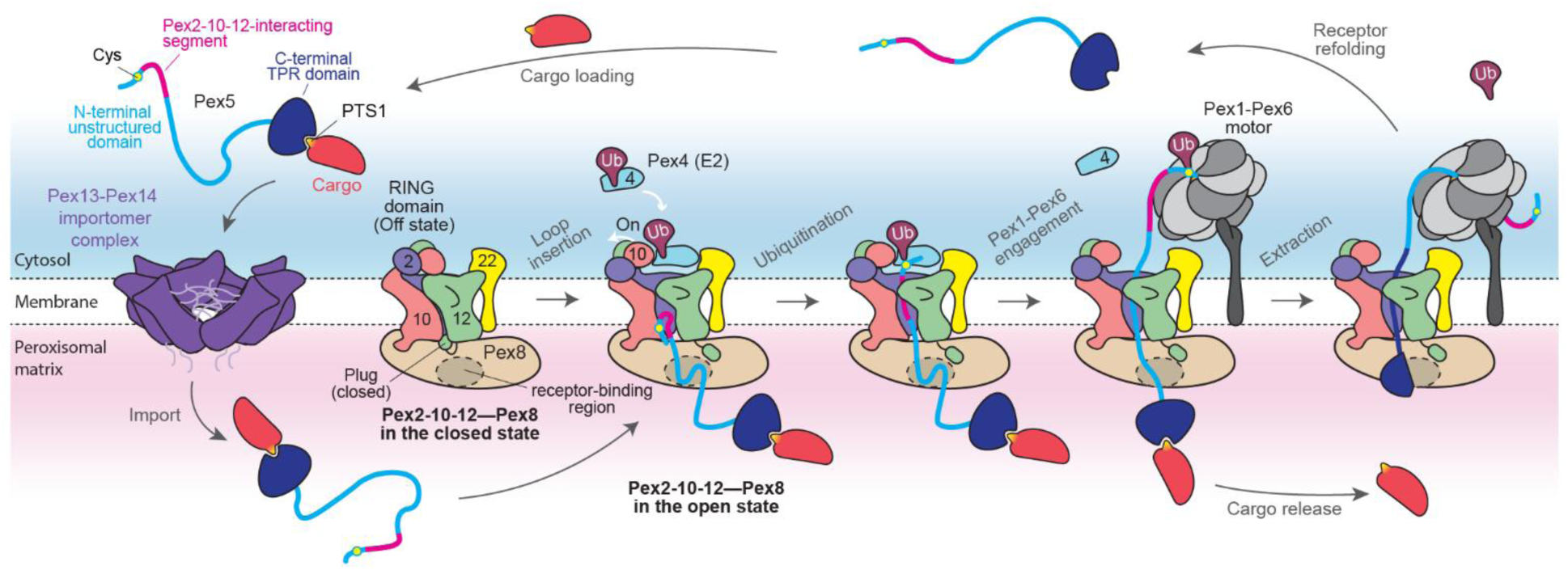
Model for receptor Pex2-10-12-mediated retro-translocation. See text.

As the insertion advances, the extreme N-terminus slips into the cytosolic funnel of Pex2-10-12. This positions Cys6 for mono-ubiquitination by Pex4 E2, which transfers the ubiquitin C-terminus onto the thiol side chain of Cys6. Proper positioning of Cys6 appears to additionally rely on interaction between the downstream segment (residues ∼11–35) of Pex5 and specific regions in Pex2-10-12. The mono-ubiquitinated N-terminus is then recognized by the Pex1–Pex6 AAA+ ATPase complex, which initiates Pex5 extraction. During this process, the C-terminal TPR domain of Pex5 is unfolded, releasing the bound cargo. Extraction likely proceeds mainly through the lateral slit beneath Pex12-TM5, whose positional flexibility may transiently widen the opening (∼15 Å) to accommodate even partially unfolded regions of Pex5.

Although the overall mechanism appears broadly conserved across eukaryotes, species-specific variations may exist, such as the absence of a Pex8-like docking factor or a Pex22-like E2-anchoring protein, neither of which has been identified in metazoans.

## DISCUSSION

E3 ubiquitin ligases are central regulators of diverse cellular processes, modulating protein functions and conferring substrate specificity for proteasomal degradation. Of more than 600 E3 ligases in humans, roughly 10% are membrane-embedded, where they play pivotal roles in quality control of membrane and lumenal proteins within organelles. Despite their importance, the functional and mechanistic understanding of these membrane-bound E3 ligases remains limited. Among them, the Pex2-10-12 complex, together with ER-localized Hrd1 and Doa10/MARCH6, represent a universally conserved subset of membrane-bound E3 ligases. Distinctively, these ligases not only catalyze ubiquitination but also enable protein retro-translocation using their multi-pass membrane domains. While recent studies indicate that retro-translocation is not a primary function of Doa10/MARCH6 (ref. ^53^), coupling between retro-translocation and ubiquitination is essential for Pex2-10-12 and Hrd1 (ref. ^54,55^). However, the molecular mechanisms coordinating these processes have been largely enigmatic.

In this study, our cryo-EM structures of the yeast Pex2-10-12 complex revealed that it does not simply form a constitutively open retro-translocation pore. Instead, the pore is gated, and this gating is coordinated with the conformational state of the RING domain such that E2∼Ub engages the RING domain for substrate ubiquitination only in the open state. Such gating likely provides two key advantages: (1) reducing off-target ubiquitination of non-substrate proteins in the closed, idle state and (2) minimizing leakage of metabolites through the pore. Our experiments using plug-deleted Pex2-10-12 as a constitutively open channel, with IPAID as a model permeant, support the latter role, while future studies will be required to address the former.

What triggers gating of the Pex2-10-12 complex remains to be resolved. One possibility is that Pex8 binding induces a conformational switch, since Pex8 interacts with both Pex10 and Pex12 and could, in principle, alter their spacing. However, this explanation seems unlikely, given that disruption of the Pex8–Pex12 interaction (Interface 2) did not significantly affect receptor recycling. A more plausible scenario is that the initial interaction between the N-terminal region of the import receptor and the lateral seam of Pex2-10-12 triggers gate opening. This would be analogous to the Sec61-mediated co-translational protein translocation, where the N-terminal signal sequence of client proteins is thought to open the lateral gate of Sec61 for insertion. In our cryo-EM analysis, import receptors were not included, yet approximately half the particle population adopted the open conformation. While detergent effects cannot be ruled out, this may also reflect a relatively modest thermodynamic energy barrier between open and closed states, which could be overcome by receptor binding. A similar substrate-independent open conformation has been observed with an archaeal Sec61 homolog^56^.

Despite architectural similarity, our yeast Pex2-10-12 structures differ in key aspects from the previously reported *Tt*Pex2-10-12 structure^23^. The *Tt* complex was captured in a single conformation that resembles the closed state of the yeast complex in terms of TM5 helix arrangement and RING domain accessibility. However, unlike the closed yeast complex, the *Tt* structure lacked a plug and displayed an open lateral seam, features more consistent with the open state of the yeast complex. The reason for this apparent hybrid conformation is unclear but may relate to the affinity tag fused to the *Tt*Pex12 N-terminus, the site that normally forms the plug. Another important difference lies in the positions of the RING domains. In *Tt*Pex2-10-12, the E2-binding surface of the Pex10-RF domain is packed against the Pex10 TMD, rendering it inaccessible to E2∼Ub, while the Pex2-RF sits directly above the cytosolic funnel, leading to the interpretation that Pex2-RF mediates receptor mono-ubiquitination. By contrast, in the yeast complex, Pex10-RF consistently occupies the funnel position regardless of the open or closed state, while Pex2-RF is oriented away from the retro-translocation path. The closed-to-open transition of the yeast complex involves only a shift in the RING domain to allow E2∼Ub access. Based on conserved structural features of RING domains, we propose that in the active state, *Tt*Pex2-10-12 also requires repositioning of Pex10-RF above the funnel, similar to the yeast complex, to enable E2∼Ub binding.

Our study identifies the function of Pex8, a peroxin of previously unknown role, as a chaperone that guides the N-termini of import receptors to the retro-translocation pore. Functional analyses show that Pex8 engages both Pex2-10-12 and Pex5, and that these interactions are indispensable for efficient peroxisomal cargo import. While this clarifies the long-standing question of why Pex8 is required, the evolutionary distribution of Pex8 raises intriguing possibilities. Pex8 is largely restricted to fungi, with no clear homolog identified in metazoans. Yet, a recent study in *Arabidopsis thaliana* uncovered a structural analog of fungal Pex8 that is crucial for peroxisomal import despite lacking obvious sequence similarity^57^. This suggests that Pex8-like mechanisms could be conserved beyond the fungal group and raises the possibility that metazoans may also harbor an as-yet-unidentified chaperone with analogous function. An alternative possibility is that metazoan Pex5 itself may directly engage with Pex2-10-12. In this scenario, the second amphipathic helix (AH2) in metazoan Pex5, located near residues 80–100, is a likely candidate for mediating this interaction, as a mutation in AH2 abolishes co-purification of the ligase^29^. Intriguingly, AH2 is specific to metazoan Pex5 and absent in yeast, suggesting that higher eukaryotes might have rewired the interaction mechanism between the receptor and Pex2-10-12. However, because metazoan Pex2-10-12 does not display clear structural features for such direct receptor binding, future studies will be needed to test this possibility.

Lastly, our study also provides mechanistic insight into how the Pex5 N-terminus inserts into Pex2-10-12. We systematically analyzed a panel of Pex5 mutants using three complementary readouts—peroxisomal import, RADAR-mediated degradation, and ubiquitination pattern—to distinguish between defects in pore insertion and mono-ubiquitination. Mutants displaying import defects coupled with increased instability likely retain pore-insertion capability but fail in efficient mono-ubiquitination, whereas those with severe import defects but normal expression and no ubiquitination likely fail in pore insertion. Two variants fit the latter category: the 5×D fusion at the N-terminus and the Δ25–35/L13S/L16S/L36S mutant. The former likely blocks insertion by raising the energetic barrier for the N-terminal slip-in upon loop insertion, while the latter likely disrupts key hydrophobic contacts required for initial insertion. Interestingly, our results reveal a degree of flexibility and redundancy in this mechanism. This may explain why scrambling residues 15–25 or introducing short deletions in this region still permits insertion, and why no strict sequence conservation is observed across species. A previous study of *Xenopus* Pex5 proposed that the segment corresponding to yeast residues 12–29 forms an amphipathic helix^29^. Mutation of three hydrophobic residues in *Xenopus* Pex5 (L17, L10, and F24; equivalent to yeast L13, L16, and T20) abolished import, although the mutant retained pore-insertion capability, likely due to residual hydrophobic residues (e.g., M18, L30, and L35) that supported insertion. Taken together, these findings suggest that the ∼20–30-residue segment immediately following the mono-ubiquitination Cys site performs two successive functions in retro-translocation: initiating loop insertion into the Pex2-10-12 pore and positioning the N-terminus of Pex5 within the cytosolic funnel for efficient mono-ubiquitination. Although the precise contribution of individual residues remains to be determined, our data strongly suggest that hydrophobic interactions between this segment and Pex2-10-12 underlie these critical steps in receptor recycling.

## Acknowledgments

This work was supported by grants from NIGMS (R01GM147628 to E.P. and training grants T32GM008295 which supported L.W., and T32GM007232 and T32GM148378 which supported N.D.), Pew Charitable Trust (Pew Biomedical Scholarship to E.P.), Shurl and Kay Curci Foundation (to N.D.), and National Science Foundation (DGE2146752 to L.W.).

## Author contributions

N.D. and L.W. performed experiments. K.G., K.Z., and J.C. assisted with experiments under supervision by N.D. and L.W. N.D. and E.P. built the atomic models of Pex2-10-12. E.P. supervised the project and wrote the manuscript with input from all authors.

## Competing interests

The authors declare no competing interests.

## Correspondence

Correspondence and requests for materials should be addressed to Eunyong Park.

## METHODS AND MATERIALS

### Yeast and *E. coli* growth Media

For routine strain maintenance and transformation, yeast cells were grown in YPD medium (1% yeast extract, 2% peptone, 2% [w/v] glucose; with additional 2% [w/v] agar in the case of solid media). YPD was supplemented with the appropriate antibiotics (500 µg/mL of hygromycin B, 600 µg/mL G418, or 100 µg/mL nourseothricin) as needed. For auxotrophic selection of yeast, drop-out media were composed of 5 g/L of ammonium sulfate, 1.7 g/L of yeast nitrogen base (United States Biological, Cat# 2030), 2% glucose, and 1.54 g/L of a drop-out mixture lacking leucine and uracil (United States Biological, Cat# D9539) or 1.92 g/L of a mixture lacking uracil (United States Biological, Cat# D9535). For −Leu drop-out, 85.6 mg/L of uracil was supplemented to the −Leu/−Ura drop-out medium. In experiments where peroxisomal protein import was tested, yeast cells were grown in an oleic acid medium consisting of 1% yeast extract, 2% peptone, 5% (v/v) glycerol, 1% (v/v) oleic acid, 0.05% (v/v) Tween-40, 0.1% or 0.2% (w/v) glucose. In all experiments, yeast cultures were grown at 30°C.

*E. coli* were grown in LB broth (1% tryptone, 0.5% yeast extract, 5g/L NaCl) supplemented with 100 µg/mL ampicillin, 34 µg/mL chloramphenicol, or 50 µg/mL kanamycin.

### Construction of plasmids and yeast strains

A list of plasmids used in this study is provided in Supplementary Table 2, yeast strains in Supplementary Table 3, and primers in Supplementary Table 4.

DNA encoding Pex5, Pex8, and Pex10 were amplified by overlapping PCR using primers (Pex5: Pex5-YTK1-gDNA1-F, Pex5-YTK1-gDNA1-R, Pex5-YTK1-gDNA2-F, Pex5-YTK1-gDNA2-R, Pex5-YTK1-gDNA3-F, and Pex5-YTK1-gDNA3-R; Pex8: pex8-f1-YTK1 and pex8-r1-YTK1, pex8-f2-YTK1 and pex8-r2-YTK1; Pex10: Pex10-YTK1-gDNA-F and Pex10-YTK1-gDNA-R) and the genomic DNA of the BY4741 yeast strain as a template and inserted into the pYTK1 entry vector of the Yeast Tool Kit system (YTK) as a Type-3 part^59^. For Pex2 and Pex12 expression, coding sequences (CDSs) lacking internal *Bsa*I and *Bsm*BI recognition sites were synthesized and inserted into pYTK1 by Gibson assembly. Where indicated, pYTK1-Pex8, pYTK1-Pex10, and pYTK1-Pex12 contain an N-terminal ALFA-tag (SRLEEELRRRLTEGS), C-terminal ALFA-tag (GSGSRLEEELRRRLTE) and C-terminal cleavable SPOT-tag (GS<u>LEVLFQGP</u>TASGPDRVRAVSHWSSGGGSGGGSTPDRVRAVSHWSSGS; underlined, an HRV 3C protease cleavage sequence), respectively, as part of the CDS. These were introduced at the position immediately after the start codon (for N-terminal tagging) or before the stop codon (for C-terminal tagging) by PCR. Additional epitope tags or fusion sequences used in this study were constructed as Type-4a parts in pYTK1. These include an HA-tag (<u>GS</u>GGGSYPYDVPDYAGSYPYDVPDYAGS; underlined, *Bam*HI site of Type 4a), ALFA-tag (<u>GS</u>GSRLEEELRRRLTE), and PTS1 (<u>GSG</u>LGRGRRSKL)^35^. Other part plasmids directly used from the original kit include pYTK18 (*ALD6* promoter), pYTK13 (*TEF1* promoter), pYTK30 (*GAL1* promoter), pYTK51 (*ENO1* terminator), pYTK61 (*ENO1* terminator), and pYTK96 (pre-assembled *URA3* integration vector).

The yeast strain (yLW006) overexpressing Pex2, Pex10, and Pex12-SPOT was generated by chromosomal integration of the YTK multigene expression plasmid (pYTK96-pGAL1-Pex2-10-12-SPOT) into the yMLT62 yeast strain. yMTL62 expresses a chimeric transcriptional activator that induces expression under the pGAL1 promoter upon addition of β-estradiol into the culture. First, CDSs for individual subunits (in pYTK1) were assembled into single-gene-expressing pYTK95 plasmids with YTK connector parts and parts for a *GAL1* promoter and an *ENO1* terminator using BsaI Golden Gate cloning. The YTK95 plasmids were then assembled into a multigene-expression plasmid (pYTK96-pGAL1-Pex2-10-12-SPOT) with pYTK96 via *Bsm*BI Golden Gate cloning. This plasmid was linearized with *Not*I endonuclease, and transformed into the yMLT62 yeast strain. Transformants were selected on a −Ura drop-out agar plate, and chromosomal integration was confirmed by PCR on genomic DNA isolated from transformants using primers URA3-5F, URA3-5R, URA3-3F, and URA3-3R.

For peroxisomal protein import assays, chromosomal integration plasmids (pYTK Kan/HO) expressing one of the reporters were constructed as follows. (1) The mRuby-PTS1 reporter plasmid (pYTK Kan/HO pTDH3-mRuby2-PTS1) for fluorescence microscopy was assembled from pYTK Kan/HO, pYTK9 (*TDH3* promoter), pYTK34 (mRuby2), pYTK61 (*ENO1* terminator), and pYTK1-PTS1 by *Bsa*I Golden Gate cloning. (2) The Ub-Y-mRuby-PTS1 reporter plasmid (pYTK Kan/HO pTDH3-UbY-mRuby2-PTS1) for fluorescence intensity was assembled from pYTK Kan/HO, pYTK9 (*TDH3* promoter), pYTK42 (Ub-Y), pYTK46 (mRuby2), pYTK61 (*ENO1* terminator), and pYTK1-PTS1 by *Bsa*I Golden Gate cloning. These reporter plasmids were linearized with *Not*I endonuclease and transformed into the BY4741 yeast strain to generate yeast strains, yLW22 (for fluorescence microscopy) and NDY41 (for the Ub-Y-mRuby-PTS1 assay). Transformants were selected on YPD-G418 agar plates, and chromosomal integration was confirmed by PCR using primers HO-5F, HO-5R, HO-3F, HO-3R and isolated genomic DNA.

The yeast strain (yLW160) for the modified version of the PDV production assay to measure IPAID permeability through plug-deleted Pex12 was created as described. First, a DNA segment for the VioA/B-expressing transcription units was amplified by PCR using primers VioA-B-YTK1-F and VioA-B-YTK1-R, and plasmid pWCD1443 as the template. The DNA segment was inserted into pYTK1 via *Bsm*BI Golden Gate cloning, generating a new Type-234 part. This VioA-VioB part was then used for *Bsa*I Golden Gate assembly with additional YTK parts (pYTK2, pYTK67, and pYTK95), to generate pYTK95-pTEF1-VioA-pTDH3-VioB. Similarly a CDS encoding VioE-mVenus-PTS1 was amplified by PCR using primers VioE-YTK1-F and VioE-YTK1-R and from plasmid pWCD2430 as the template and inserted in pYTK1 as a new Type-3b part. The VioE-mVenus-PTS1 CDS was assembled together with additional YTK parts (pYTK3, pYTK13 [*TEF1* promoter], pYTK43 [Ub-R], pYTK51 [*ENO1* terminator], pYTK68, and pYTK95) via *Bsa*I Golden Gate cloning, resulting in plasmid pYTK95 pTEF1-Ub-R-VioE-mVenus-ePTS1. The two plasmids, pYTK95 VioA-VioB and pYTK95 Ub-R-VioE-PTS1, were assembled into a multigene-expressing chromosomal integration plasmid (separately made from pYTK8, pYTK47, pYTK73, pYTK77, pYTK88, pYTK90, and pYTK94) via *BsmB*I Golden Gate cloning. The resulting plasmid, pYTK Kan/HO pTEF1-VioA-pTDH3-VioB-pTEF1-UbiR-VioE-mVenus-ePTS1, was linearized by *Not*I endonuclease and used for transformation of the BY4741 strain. G-418-resistant transformants were selected on a YPD agar plate, and integration was confirmed by PCR using primers HO-5F/HO-5R and HO-3F/HO-3R.

For deletion of the chromosomal *PEX5, PEX8, PEX10,* and *PEX12* genes in yeast, deletion cassettes were constructed by PCR amplification of the *URA3* marker from pYTK74 or the *LEU2* marker from pYTK75 using primers (PEX5: pex5d-F, pex5d-R; PEX8: pex8d-F, pex8d-R; PEX10: pex10d-F, pex10d-R; PEX12: pex12d-F, pex12d-R). The PCR products were directly used for transformation of yeast. Deletion was confirmed by PCR analysis of isolated genomic DNA using primers pex8d-5d-conf-F/pex5d-conf-R for PEX5, pex8d-5d-conf-F/pex8d-conf-R for PEX8, pex10d-conf-F/pex10d-conf-R for PEX10, and pex12d-conf-F/pex12d-conf-R for PEX12.

Functional complementation experiments were performed by chromosomal integration of a plasmid (pYTK Hyg/leu2) expressing a WT or mutant copy of Pex8, Pex10, Pex12, or Pex5 into respective *PEX* deletion strains. Unless stated otherwise, the Pex proteins were expressed under a constitutive *ALD6* promoter (from pYTK18). However, some Pex5 mutants exhibited substantially lower expression levels than that of WT, and for these mutants, we also tested a stronger constitutive promoter, *pTEF1* (from pYTK13). All plasmids were constructed using the YTK system and *Bsa*I Golden Gate cloning. Used parts were listed in Supplementary Table 3. Constructed integration plasmids were digested with *Not*I endonuclease prior to transformation. Transformants were selected on YPD-hygromycin agar plates, and chromosomal integration was confirmed by PCR using primers LEU2-5F/LEU2-5R and LEU2-3F/LEU2-3R.

For *E. coli* expression of Pex8 fused to the maltose-binding protein (MBP-Pex8), pMAL-c2x-Pex8-His_6_ was constructed by PCR amplification of a CDS for Pex8-His_6_ using primers pex8-pMAL-F and pex8-pMAL-R and pYTK1-ALFA-Pex8 as template. In the pull-down experiments in Fig. 3E, Pex8 lacks its native C-terminal PTS1 (ΔSKL), and this deletion was introduced by PCR. The PCR product was inserted into the *EcoR*I—*Pst*I sites of pMAL-c2x by standard restriction digestion and ligation. Similarly, for *E. coli* expression, a CDS for Pex5-Strep was amplified by PCR and inserted into a pQE plasmid downstream of the His-Sumo-tag.

### Protein purification

The Pex2-10-12 complex was purified from yeast strain yLW6 overexpressing Pex2, Pex10, and SPOT-tagged Pex12. Cells were grown in the oleic acid medium until they reached an OD_600_ of 0.8–1.0. Overexpression was induced by the addition of 50 nM β-estradiol for 5 hours. Cells were harvested by centrifugation, resuspended with a small volume of lysis buffer (50 mM Tris-HCl pH 8.0, 200 mM NaCl, 2 mM phenylmethylsulfonyl fluoride [PMSF], 5 µg/mL aprotinin, 25 µg/mL leupeptin, and 1 µg/mL pepstatin A), frozen in liquid nitrogen, and stored at −80°C.

Frozen cells were pulverized with a freezer mill device (Spex Sampleprep, Model 6870), and the resulting powder was mixed with additional ice-cold lysis buffer. All subsequent steps were performed at 4°C. Unbroken cells were removed by centrifugation (3,000 g, 10 min). Membrane fractions were collected by ultracentrifugation (140,000 g, 1.5 hours). The membrane pellet was resuspended in lysis buffer containing 1% lauryl maltose neopentyl glycol (LMNG; Anatrace, Cat# NG310) and 0.2% cholesteryl hemisuccinate (CHS; Anatrace, Cat# CH210), and the membranes were solubilized by gentle agitation for 1 hour. The lysate was clarified by ultracentrifugation (125,000 g, 1 hour). The supernatant was supplemented with 10% glycerol and 6 µg of Benzonase nuclease, and incubated with 2 mL of home-made BC2-nanobody conjugated sepharose resin for 2 hours. The resin was then packed in a gravity column and washed with 30 column volumes (CVs) of wash buffer (50 mM Tris HCl pH 8.0, 200 mM NaCl, 10% glycerol, and 0.04% glyco-diosgenin [GDN; Anatrace, Cat# GDN101]). Proteins were eluted by cleaving the SPOT-tag by incubating the resin overnight with human rhinovirus (HRV) 3C protease. The eluate was collected and concentrated to 500 µL with an Amicon concentrator (Cytiva, 100-kDa cut-off) and injected to a Superose 6 Increase 10/300 GL column (Cytiva) equilibrated with 50 mM Tris-HCl pH 8.0, 200 mM NaCl, and 0.04% GDN.

MBP-Pex8-His_6_ was purified recombinantly from *E. coli*. The pMAL-c2x Pex8-His_6_ plasmid was introduced into BL21(DE3) cells by transformation. Cells were grown in LB medium at 37°C until reaching an OD_600_ of 0.6–0.8. MBP-Pex8-His_6_ overexpression was induced by addition of 0.5 mM Isopropyl-β-D-thiogalactopyranoside (IPTG), and cultures were incubated at 22°C for 6 hours or overnight at 18°C before harvesting. Cell pellets were frozen in liquid nitrogen and stored at −80°C until use. All subsequent steps were performed at 4°C. The cell pellet was thawed and resuspended in lysis buffer (50 mM Tris-HCl pH 8.5, 300 mM NaCl, 10% glycerol, 10 mM imidazole, 2 mM PMSF, 5 µg/mL aprotinin, 25 µg/mL leupeptin, and 1 µg/mL pepstatin A) and lysed by sonication. The lysate was clarified by ultracentrifugation (125,000 g, 1 hour). The supernatant was then supplemented with 12 µg of Benzonase nuclease and incubated with 3 mL of His-Pur cobalt resin (Thermo Scientific; Cat# 89965) for 2 hours. The resin was washed with 30 CVs of wash buffer (50 mM Tris-HCl pH 8.5, 300 mM NaCl, 10% glycerol, and 25 mM imidazole), and protein was eluted with 5 CVs of elution buffer (50 mM Tris-HCl pH 8.5, 300 mM NaCl, 10% glycerol, and 200 mM imidazole). The eluate was supplemented with 1 mM ethylenediaminetetraacetic acid (EDTA) and 2 mM dithiothreitol (DTT) and dialyzed overnight against dialysis buffer (50 mM Tris-HCl pH 8.5, 300 mM NaCl, 10% glycerol, 1 mM EDTA, and 2 mM DTT) using dialysis tubing with a 10-kDa molecular weight cut-off. After concentrating to 500 µL using an Amicon concentrator (50-kDa cut-off), the sample was injected into a Superdex 200 Increase 10/300 GL column (Cytiva) equilibrated with 50 mM Tris-HCl pH 8.5, 200 mM NaCl. For biochemistry experiments, protein was supplemented with 10% glycerol, concentrated to 1 mg/mL, frozen in liquid nitrogen, and stored at -80°C.

C-terminally Strep-tagged Pex5 protein was recombinantly purified from *E. coli*. BL21(DE3) cells were transformed with the pQE-His-SUMO-Pex5-Strep plasmid and grown in LB medium at 37°C until reaching an OD_600_ of 0.6–0.8. Overexpression was induced with 0.5 mM IPTG at 18°C for ∼20 hours. Cells were resuspended in lysis buffer (50 mM Tris-HCl pH 8.0, 500 mM NaCl, 1 mM MgCl_2_, 25 mM imidazole, 2mM PMSF, 5 µg/mL aprotinin, 25 µg/mL leupeptin, and 1 µg/mL pepstatin A) and lysed by sonication. After clarifying by centrifugation, the lysate was incubated with His-Pur cobalt resin, and the resin was washed with 20 CV of wash buffer (50 mM Tris-HCl pH 8.0, 500 mM NaCl, 1 mM MgCl_2_, 50 mM imidazole, and 10% glycerol) and eluted with 3 CV of elution buffer (50 mM Tris-HCl pH 7.5, 500 mM NaCl, and 300 mM imidazole). The eluate was dialyzed against 50 mM Tris-HCl pH 8.0 and 500 mM NaCl). The His-SUMO-tag was cleaved by addition of the Ulp1 protease, and the cleaved His-SUMO fragments were removed by flowing the sample through His-Pur cobalt resin. 1 mM EDTA and 2 mM DTT were added to Pex5-Strep, and the sample was concentrated and injected to a Superdex 200 column equilibrated with buffer containing 50 mM Tris-HCl pH 8.0, 500 mM NaCl, 1 mM EDTA, and 2 mM DTT. Peak fractions were pooled, supplemented with 10% glycerol, concentrated to ∼1 mg/mL, frozen in liquid nitrogen, and stored at −80°C until use.

### Cryo-EM analysis

Two datasets were collected for the Pex2-10-12 complex bound to MBP-Pex8. Slightly different sample preparation methods were used for the two datasets, but resulting structures showed no discernible differences. Purified Pex2-10-12 and MBP-Pex8-His_6_ were mixed at molar ratios of 1:1.2 for Dataset 1, or 1:1.5 for Dataset 2, and concentrated to ∼6 mg/mL (Dataset 1) or ∼12 mg/mL (Dataset 2) using an Amicon concentrator (100-kDa cut-off). Samples were supplemented with 3 mM fluorinated-fos-choline-8 (Anatrace, Cat#F300F). 3µL of sample was applied to a gold holey carbon grid (Quantifoil R 1.2/1.3, 400 mesh), which was glow-discharged using a PELCO-easiGlow (0.39 mBar, 25–30 mA, 40 s), and plunge frozen in liquid ethane using a Vitrobot Mark IV (FEI). Blotting was performed with Whatman No. 1 filter papers at 4°C with ∼100% relative humidity.

Both datasets were collected on a Krios G3i microscope (Thermo Fisher FEI), operating at 300 keV, equipped with a Gatan Quantum Energy Filter (slit width of 20 eV), and a K3 direct electron detector (Gatan). Images were collected as dose-fractionated movies (50 electrons per Å^2^ applied over 50 frames) at a physical pixel size of 1.05 Å and a target defocus range of 0.8–1.6 µm. Microscope control and data acquisition were performed using SerialEM^60^ with image-shift multi-shots and coma correction via beam tilt compensation.

The cryo-EM data processing procedure is summarized in Supplementary Fig. 1F. In brief, initial motion correction, contrast transfer function (CTF) estimations, and automatic particle picking were performed in Warp^61^. Particles were imported into CryoSPARCv4 (ref. ^62^) and 2D classification and Ab-initio reconstruction were performed. The resulting 3D model was used to generate templates for template picking in CryoSPARC. The newly picked particle set was subjected to 2D classification and Ab initio reconstruction. The particle set was further cleaned by Heterogeneous refinement, and a high-resolution consensus map was obtained using Non-uniform refinement. The particles for this map were then used to train the Topaz particle picker^63^. Particles were re-picked from motion-corrected micrographs using Topaz and subjected to rounds of 2D classification, Ab-Initio reconstruction, Heterogeneous refinement, Non-uniform refinement, and Local CTF refinement. Datasets 1 and 2 were processed separately using similar workflows and then selected particles were combined at this step. The combined particle set was applied to 3D classification and split into the two visually distinct classes, deemed ‘open’ and ‘closed’ conformations. Each group was subjected to an additional round of Heterogeneous refinement and a final round of Non-uniform refinement. Maps were B-factor sharpened and filtered according to the default Non-uniform refinement function and used for atomic model building and refinement. For images shown in Figures 1 and 4, the density maps were sharpened using DeepEMhancer using its default tight-target model^64^.

### Cryo-EM atomic model building

Initial atomic models of the Pex2-10-12–Pex8 complex were generated by docking an AlphaFold3 model^65^ into the cryo-EM maps. Models were rebuilt manually using local refinement in Coot^66^. In the closed-state model, a RING domain model generated by AlphaFold3 was docked into the unsharpened cryo-EM map as a rigid body, and no additional model refinement was performed in Coot due to limited resolution in this region. Several regions, including the plug domain and TM5 helices, were built de novo as its cryo-EM map features differed from the AlphaFold model (commonly low-pLDDT regions). Global refinement was performed using Phenix (phenix.real_space_refine) with resolution set to the nominal resolution of the cryo-EM maps^67^. The models were validated using MolProbity^68^. Figures were prepared using PyMOL (Schrödinger, Inc.), UCSF Chimera^69^, and UCSF ChimeraX^70^.

### Sequence alignments

Amino acid sequences of Pex2, Pex10, Pex12, and Pex5 from fungal species were retrieved from NCBI’s ClusteredNR database using BLASTp queries of respective *S. cerevisiae* sequences (full length sequences for Pex2, Pex10, and Pex12; residues 1–300 for Pex5). Small subsets of sequences were removed using a sequence length filter (200–510 for Pex2 and Pex10, and 300–510 for Pex12). Sequences were aligned using MAFFT on the EMBL-EBI web server. Amino acid conservation was mapped onto the closed-state cryo-EM structure using Chimera. For Pex5, the sequence logo representation was generated using WebLogo 2.8.2.

### Confocal fluorescence microscopy

Confocal microscopy was performed with a Zeiss Axio Observer Z1 microscope equipped with a spinning disk confocal unit (CrestOptics, X-Light V3), a laser light source (89 North, LDI-7), a 100x/1.4NA oil-immersion objective (Zeiss, Plan-Apochromat 100x/1.4 Oil), and an sCMOS camera (PCO, pco.edge 4.2bi). Yeast strains carrying a peroxisome reporter mRuby-PTS1 were inoculated into the oleic acid liquid medium supplemented with 0.1% glucose and grown for 1 day at 30°C. Cells were washed in water and resuspended in live cell imaging solution (Invitrogen, Cat# A14291DJ), applied to a #1.5 glass coverslip, and mounted to slides. A 555-nm laser was used for excitation and mRuby signals were acquired over three Z-steps with an interval of 0.5 um and an exposure time of 500-ms. For data presentation, the z stacks were combined into a maximum intensity projection using FIJI^71^. Brightness was adjusted uniformly across all conditions with FIJI, and overlaid DIC images were used to demarcate cell boundaries (DIC images were removed for clarity).

### Ub-Y-mRuby-PTS1 reporter assay

A single colony of a given strain was picked and inoculated in the oleic acid medium as in confocal microscopy experiments. In Figs. 1–5, the medium was supplemented with 0.1% glucose, whereas in Fig. 6, the medium was supplemented with 0.2% glucose. Cells were grown for ∼16 hours at 30°C, harvested, and resuspended to an OD_600_ of 1.0 in buffer containing 50 mM Tris-HCl pH 7.5 and 150 mM NaCl. 100 µL of the cell suspension were transferred to a black 96-well microplate, and fluorescence intensities at 615 nm (565 nm for excitation) were measured using a plate reader (BMG Labtech, ClarioStar). Intensity values were blank corrected and normalized with respect to the value from a reference strain (WT non-deletion parent) measured at the same time. Relative fluorescence intensities with respect to the strain transformed with the WT-expressing plasmid were plotted using RStudio software with the ggplot2 package. WT and deletion values for given peroxins are reused across figures.

### Violacein intermediate-production assay

Yeast were grown overnight in the YPD medium, and cultures were diluted to an OD_600_ of 0.05. 5 uL were spotted onto a synthetic complete agar plate (0.17% YNB, 0.5% ammonium sulfate, 0.2% synthetic complete amino acid mixture [US biological, Cat# D9515], 2% glucose, 2% bacto-agar, and 400 ug/mL G418). Plates were incubated for 3 days at 30°C plates before taking images.

To quantify PDV levels in the permeability assay, yeast cells were collected from the agar plate and resuspended in water. 1.5 OD of cells were spun down and resuspended in 150 uL acetic acid. The cell suspension was then heated at 95°C for 30 min. After removing insoluble debris by centrifugation (17,000 g, 10 min), 100uL of the supernatant was collected, and the fluorescence intensity at 585 nm upon excitation at 530 nm was measured using a plate reader (BMG Labtech, Clariostar). Intensity values from *pex12Δ* cells were used as the background. Background-substracted values were then normalized with respect to values from the strain with the WT Pex12-expressing plasmid.

### *In-vitro* pull-down assays

For the pull-down experiments of MBP-Pex8 by the Pex2-10-12 complex, Pex2-10-12-SPOT was immobilized onto BC2T nanobody resin as described above, but without the HRC 3C protease cleavage step. 100 µL of the resin slurry was incubated with MBP-Pex8-His_6_ (WT or indicated mutant) for 30 min at 4°C, washed five times with 1 mL of buffer containing 50 mM Tris-HCl pH 8.0, 200 mM NaCl, 1 mM EDTA, 2 mM DTT, 10% glycerol, and 0.04% GDN. Proteins were eluted by addition of SDS sample buffer to beads and incubation at 75°C for 5 min. The eluates were analyzed by SDS-PAGE and Coomassie staining.

For the pull-down experiments of Pex8 by Pex5, purified full-length WT Pex5-Strep was mixed with either WT, mut3 (F335D/I288D), or mut3/4 (L479N/I435N/T432R/F335D/I288D) MBP-Pex8_ΔSKL_-His_6_ at equimolar (1 uM) concentrations and incubated for 30 min on ice. The mixtures were incubated with 20 uL of Strep-Tactin XT agarose beads (IBA, Cat# 2-5030-002) for 30 min at 4°C. The beads were washed five times with 1 mL buffer containing 50 mM Tris-HCl pH 8.5, 200 mM NaCl, and 10% glycerol. Proteins were eluted by incubating beads with 40 uL of buffer additionally containing 50 mM biotin. The eluates were analyzed by SDS-PAGE and Coomassie staining.

### Pex5 ubiquitination assay

Yeast starter cultures grown in the oleic acid medium supplemented with 0.1% glucose were inoculated into fresh medium, and cells were grown for an additional 16 hours. Equal numbers of cells were harvested by centrifugation, washed once with water, and resuspended in 700 µL of 5% trichloroacetic acid (TCA). Cells were lysed by beating with glass beads. Precipitated proteins were collected by centrifugation, washed twice with ice-cold acetone, and air dried. Proteins were resuspended with urea SDS buffer (50mM Tris-HCl pH 7.5, 6 M Urea, and 1% SDS) and heated at 75°C for 10 min. Insoluble matter was removed by centrifugation, and the supernatant was diluted 10-fold with native buffer containing 50 mM Tris-HCl pH 7.5, 150 mM NaCl, 0.5% Tween-20, 2 mM EDTA, 2 mM PMSF, 25 mM N-ethylmaleimide, 5 µg/mL aprotinin, 25 µg/mL leupeptin, and 1 µg/mL pepstatin A. Lysates were then incubated with anti-HA affinity resin (Abcam, Cat# ab270603) for 1 h at 4°C. The resin was washed three times with 1 mL native buffer, and proteins were eluted with an SDS sample buffer containing 6 M urea but lacking β-mercaptoethanol (β-ME). Samples were heated for 10 min at 42°C before SDS-PAGE analysis. For β-ME-treated samples, 3.3% β-ME was added before heating.

### SDS-PAGE and Immunoblotting

For expression verification of mutant Pex proteins, yeast cells were grown overnight in the oleic acid medium. 0.5 OD of cells were harvested by centrifugation, resuspended in 700 µL 10% TCA, and incubated on ice for 30 min. Cells were spun down, washed twice with ice-cold acetone, and air dried. The pellet was resuspended in 150 µL urea buffer (50 mM Tris-HCl pH 7.5, 8 M Urea, 5 mM EDTA, and 2% [v/v] β-ME) and beaten with glass beads. The lysate was recovered, mixed with 37.5 µL 5x SDS sample buffer, and heated for 20 min at 42°C. After clarification by centrifugation (17,000 g, 10 min), 15 µL of supernatant were analyzed by SDS-PAGE and immunoblotting.

All SDS-PAGE were performed using 10% bis-tris gels. with the exception of Fig. 1A, where a 10% tris-glycine gel was used.

The following antibodies were used for immunoblots, rabbit α-HA-tag (Proteintech, Cat# 51064-2-AP; 1:5,000) or mouse α-HA-tag (Invitrogen, Cat# 26183; 1:4,000), goat α-mouse-IgG-HRP (Thermo Fisher, Cat# 31430; 1:10,000), and goat α-rabbit-IgG-HRP (Thermo Fisher, Cat# 31460). ALFA-tagged and SPOT-tagged proteins were detected with homemade ALFA-nanobody-Fc and BC2T-nanobody-Fc as previously described^72^.

## Data Availability

The cryo-EM maps and atomic models are deposited to EM Data Bank (EMDB) and Protein Data Bank (PDB) under the following accession IDs: EMD-72627 and PDB 9Y6Q for the closed-state structure; EMD-72628 and PDB 9Y6R for the open-state structure.

## Notes

### Competing Interest Statement

The authors have declared no competing interest.

