## Supplementary Information for "Structural basis of receptor retro-translocation in peroxisomal protein import"

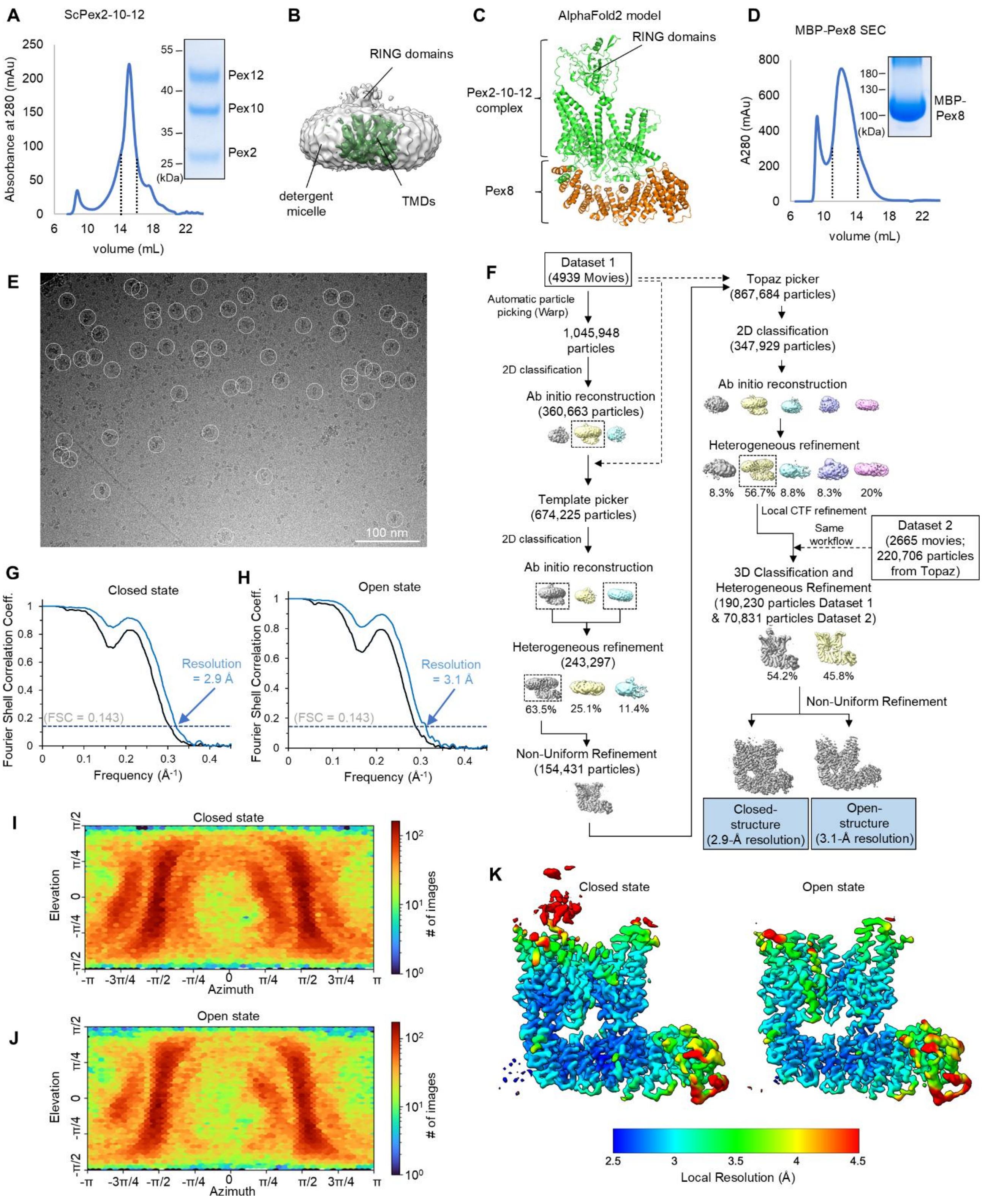

**Supplementary Figure 1. Single-particle cryo-EM analysis of the yeast Pex2-10-12 complex.**
(**A**) Purification of yeast Pex2-10-12. Overexpressed Pex2-10-12 was affinity purified by C-terminal SPOT-tagged attached to Pex12 and subjected to Superose 6 size-exclusion chromatography. The fractions between the two dotted lines were analyzed by SDS-PAGE, pooled, and used for cryo-EM. (**B**) Low-resolution cryo-EM reconstruction of yeast Pex2-10-12 without Pex8 from ~150,000 particles images obtained in a comparable imaging condition used for the high-

resolution structures in this study. (**C**) AlphaFold prediction of complex formation between yeast Pex2-10-12 and Pex8. (**D**) The MBP-Pex8 fusion protein was recombinantly expressed and purified from *E. coli* using a C-terminal His-tag and subjected to Superdex 200 size-exclusion chromatography. The fractions between the two dotted lines were analyzed by SDS-PAGE, pooled, and used for cryo-EM. (**E**) Representative cryo-EM images of Pex2-10-12–Pex8 particles (white circles). Scale bar represents 100 nm. (**F**) Flow chart summarizing the cryo-EM image processing. (**G**, **H**) Fourier shell correlation (FSC) curves for the final 3D refinements of the closed and open-state structures. Black line, spherical mask; blue line, tight-masked corrected FSC. (**I**, **J**) Particle orientation distribution for the final 3D refinements. (**K**) Local resolution distribution of the final 3D refinement maps.

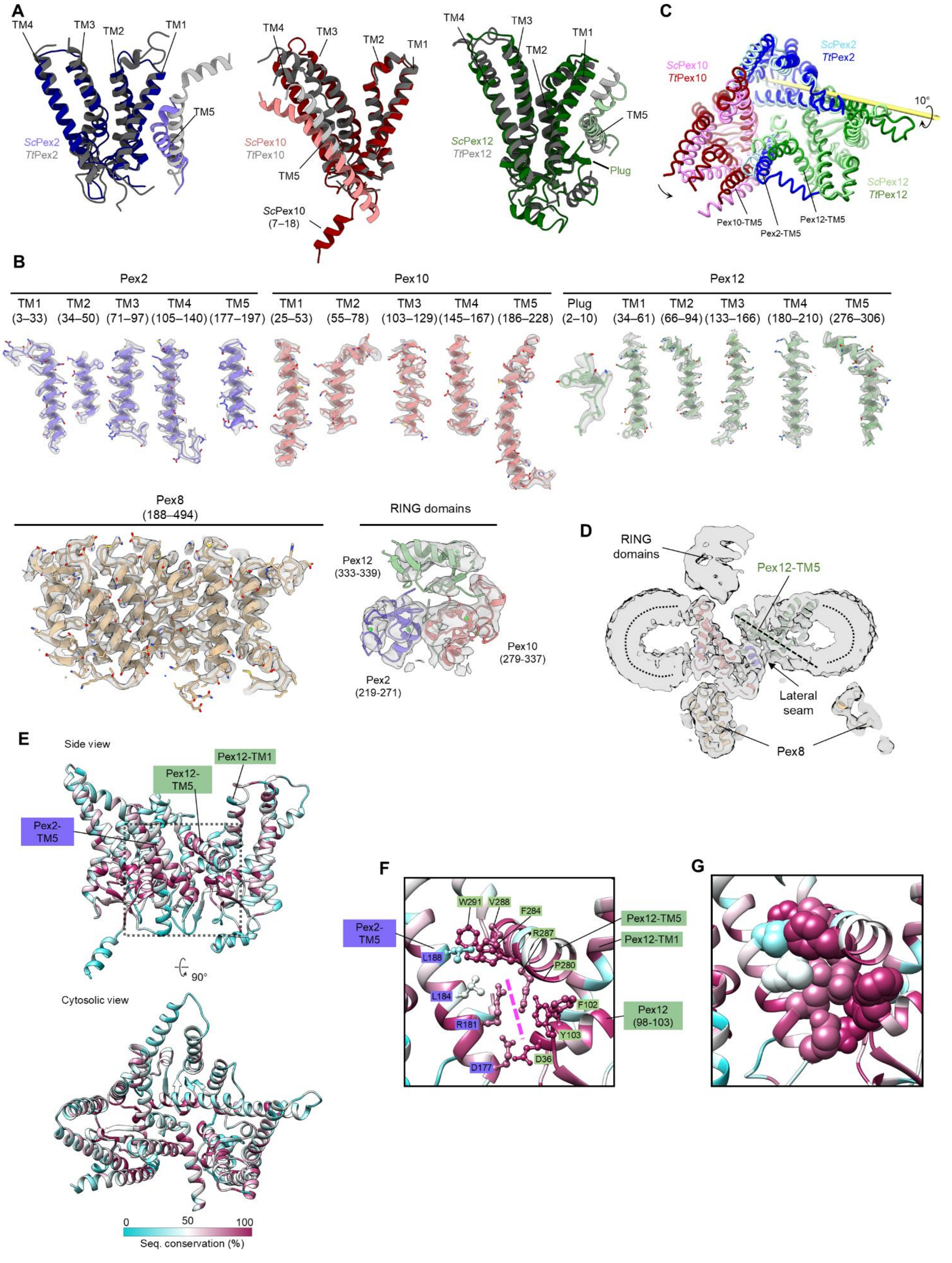
A
TM4
TM3
TM2
TM1
TM5
ScPex2
TtPex2
ScPex10
TtPex10
ScPex10
(7–18)
ScPex12
TtPex12
Plug
C
ScPex2
TtPex2
ScPex10
TtPex10
10°
ScPex12
TtPex12
Pex10-TM5
Pex2-TM5
Pex12-TM5
B
Pex2
Pex10
Pex12
TM1 (3–33)
TM2 (34–50)
TM3 (71–97)
TM4 (105–140)
TM5 (177–197)
TM1 (25–53)
TM2 (55–78)
TM3 (103–129)
TM4 (145–167)
TM5 (186–228)
Plug (2–10)
TM1 (34–61)
TM2 (66–94)
TM3 (133–166)
TM4 (180–210)
TM5 (276–306)
Pex8 (188–494)
RING domains
Pex12 (333-339)
Pex2 (219-271)
Pex10 (279-337)
D
RING domains
Pex12-TM5
Lateral seam
Pex8
E
Side view
Pex12-TM1
Pex12-TM5
Pex2-TM5
90°
Cytosolic view
0
50
100
Seq. conservation (%)
F
W291
V288
F284
R287
P280
L188
L184
R181
D177
F102
Y103
D36
Pex2-TM5
Pex12-TM5
Pex12-TM1
Pex12 (98-103)
G

**Supplementary Figure 2. Structures of the Pex2, Pex10, and Pex12 subunits and cryo-EM map quality.**
(**A**) TM arrangements of Pex2, Pex10, and Pex12. Colored ribbons show the cryo-EM structure of yeast (*Sc*) Pex2-10-12 in the closed-state, and gray ribbons show the cryo-EM structure of *Thermothelomyces thermophilus* (*Tt*) Pex2-10-12 (PDB 7T92). (**B**) Representative atomic model segments overlaid with the density map. Numbers indicate amino acid residue ranges. (**C**) Alignment of *Tt*Pex2-10-12 with *Sc*Pex2-10-12 based on Pex12. Shown is the top view from the cytosol. The yellow rod marks the rotational axis for the positional deviation of *Sc*Pex10 relative to *Tt*Pex10. (**D**) Cross-sectional view of the cryo-EM density map highlighting the tilted TM5 helix (dashed line) of Pex12 and the lateral seam beneath it. A low-pass filter was applied to the density map to visualize the detergent micelle (outlined with dotted curves). (**E**) Amino acid sequence conservation map derived from alignments of 110 Pex2, 482 Pex10, and 421 Pex12 clustered sequences from fungal species. The boxed area is shown enlarged in panel F. (**F**) Side view of the lateral seam region. The magenta dashed line marks the seam that opens in the open-state structure. (**G**) As in F, but with side chains shown as spheres to illustrate tight packing and high sequence conservation at the seam.

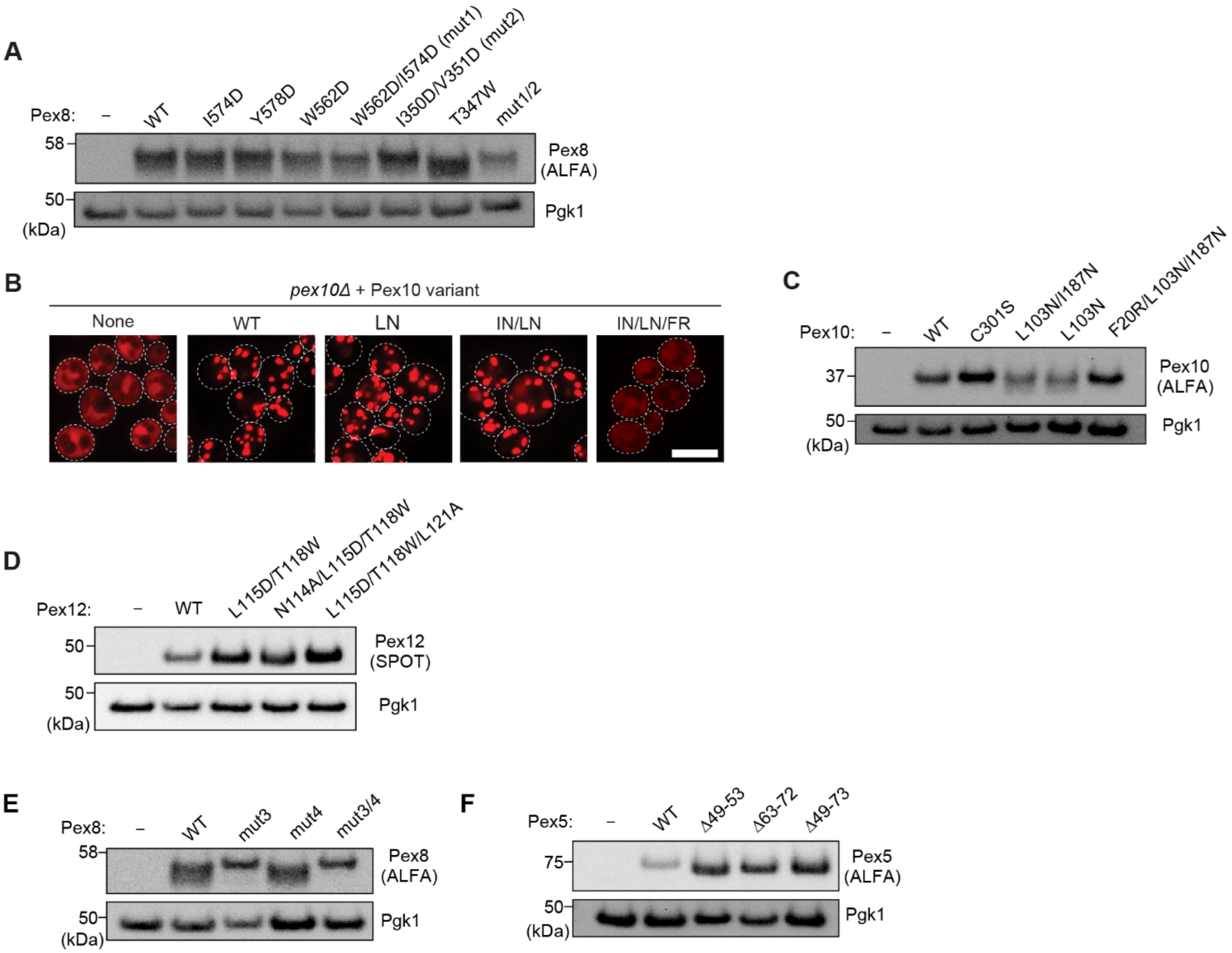

**Supplementary Figure 3. Interactions of Pex8 with Pex2-10-12 and Pex5 are essential for peroxisomal protein import in yeast.**
(**A**) Immunoblot confirming expression of N-terminally ALFA-tagged Pex8 mutants used in Fig. 2. Pgk1, loading control. (**B**) Effects of Pex10 Interface-1 mutations on peroxisomal localization of mRuby-PTS1 were examined by confocal microscopy. LN, L103N; IN, I187N; FR, F20R. Scale bar, 5 μm. (**C**, **D**) Immunoblot verifying expression of Pex10 (C) and Pex12 (D) mutants. Pex10 and Pex12 constructs contain C-terminal ALFA and SPOT-tags, respectively. (**E**, **F**) Immunoblots verifying expression of ALFA-Pex8 (E) and Pex5-ALFA (F) mutants. Pgk1, loading control.

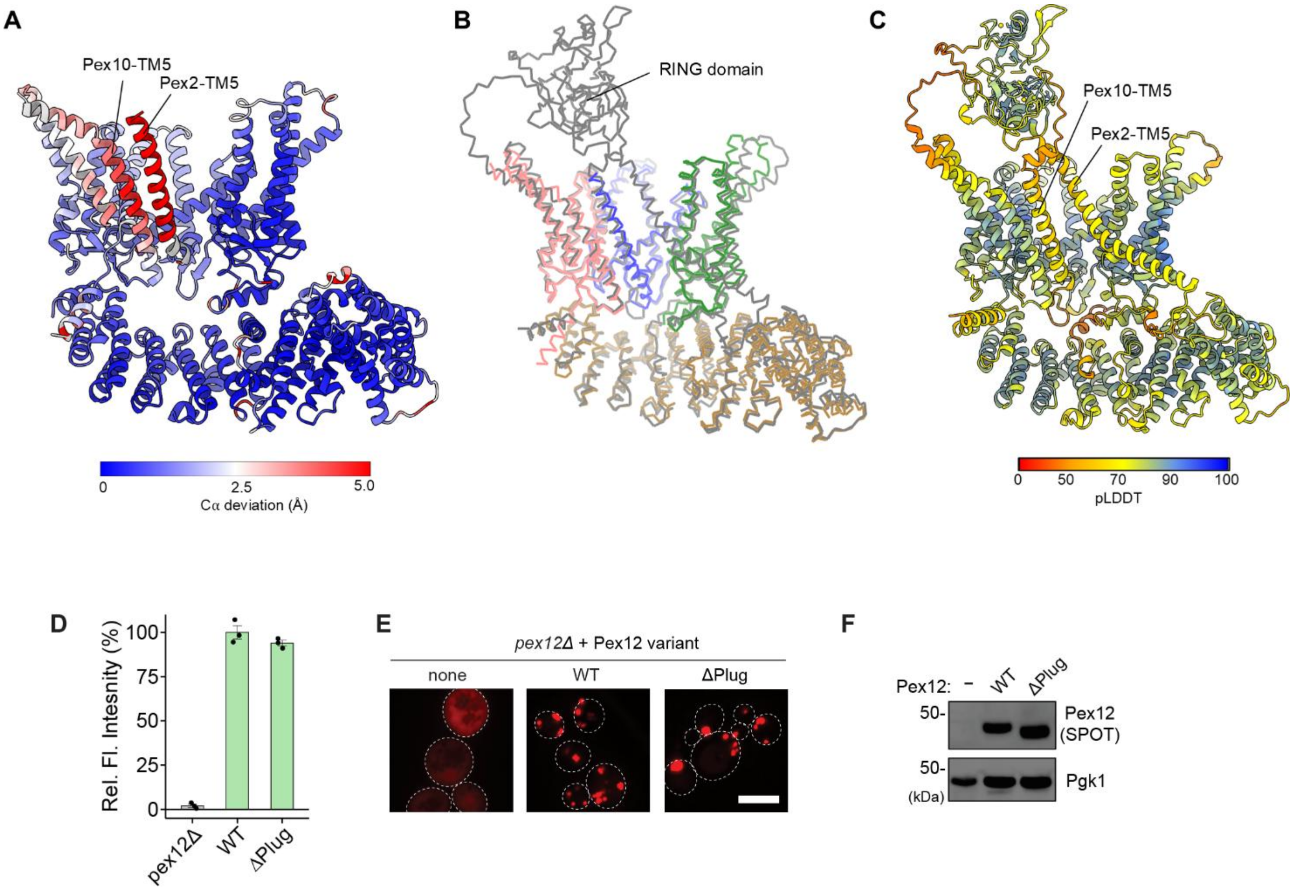

**Supplementary Figure 4. Comparison of the open-state Pex2-10-12–Pex8 cryo-EM structure with the closed-state structure and an AlphaFold model.**
(**A**) The open-state cryo-EM structure was aligned with the closed-state structure, and per-residue Cα deviations were mapped as a heat map onto the open-state structure. (**B**) Alignment of the open-state cryo-EM structure (colored) with an AlphaFold prediction for yeast Pex2-10-12–Pex8 (gray). (**C**) pLDDT confidence score map of the AlphaFold prediction in B. (**D**, **E**) The plug-deleted (ΔPlug) Pex12 mutant retains WT-like peroxisomal import activity as measured by the Ub-Y-mRuby-PTS1 reporter assay (D) and mRuby-PTS1 microscopy assay (E). In D, data represent means ± s.e.m (n=3). In E, scale bar, 5 µm. (**F**) Immunoblot confirming expression of the ΔPlug Pex12 mutant. Pex12 contains a C-terminal SPOT-tag. Pgk1, loading control.

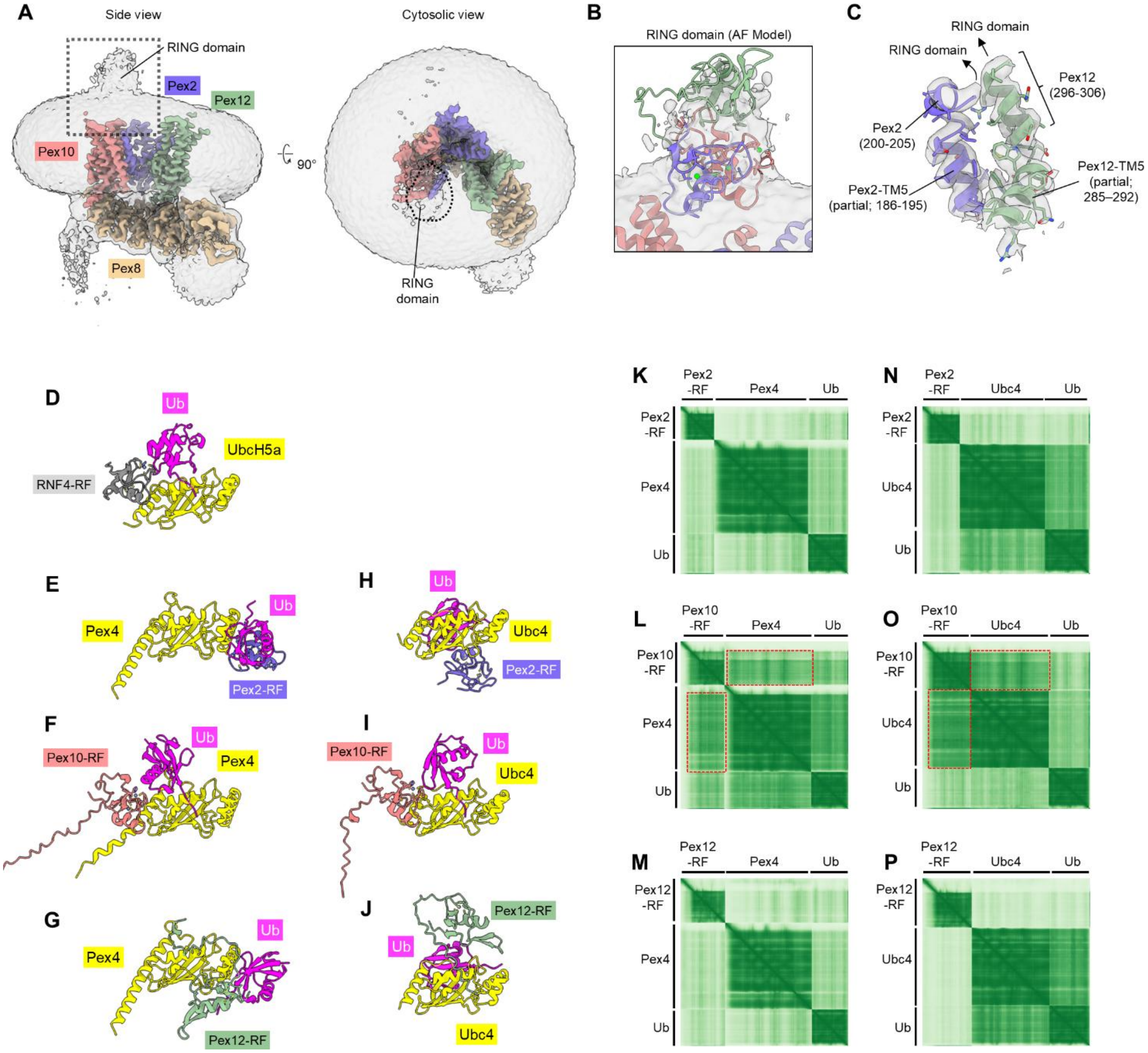

**Supplementary Figure 5. Interaction between E2 ubiquitin-conjugating enzymes and the Pex2-10-12 RING-finger domains in the open state.**
(**A**) Position of the RING domain in the open-state Pex2-10-12–Pex8 cryo-EM structure. The unsharpened density map (gray, 1.4σ contour) is overlaid with the high-resolution features (colored) of the Pex2-10-12 TMD and Pex8. Density corresponding to the RING domain is indicated. A close-up of the boxed region is shown in B. (**B**) Close-up of the RING domain feature in the open-state cryo-EM density map. The AlphaFold model (ribbon representation) of yeast Pex2-10-12–Pex8 is aligned with the open-state cryo-EM structure. (**C**) TM5 regions of Pex2 and Pex12 that position the RING domain in the closed-state structure, shown with the atomic model (ribbon) and density map (gray). In the open-state cryo-EM structure, Pex12-TM5 is flexible and invisible in the density map. (**D**) Co-crystal structure of the RNF4 RING-finger (RF) domain, UbcH5a (E2), and ubiquitin (Ub), showing canonical RF:Ub:E2 interfaces (PDB 4AP4; ref. [41]). (**E**–**G**) AlphaFold predictions of the Pex4 E2, ubiquitin, and the RF domain from Pex2 (E) or Pex10 (F), or Pex12 (G). The orientation of Pex4 is the same as that in D. (**H**–**J**) As in E–G, but with Ubc4 as the E2 enzyme instead of Pex4. (**K**–**M**) Predicted aligned error (PAE) matrices form the AlphaFold predictions in E–G. In L, the regions outlined by red dashed lines correspond to the Pex4–Pex10-RF interaction. (**N**–**P**) PAE matrices from the AlphaFold predictions in H–J. In O, the regions outlined by red dashed lines correspond to the Ubc4:Pex10-RF interaction.

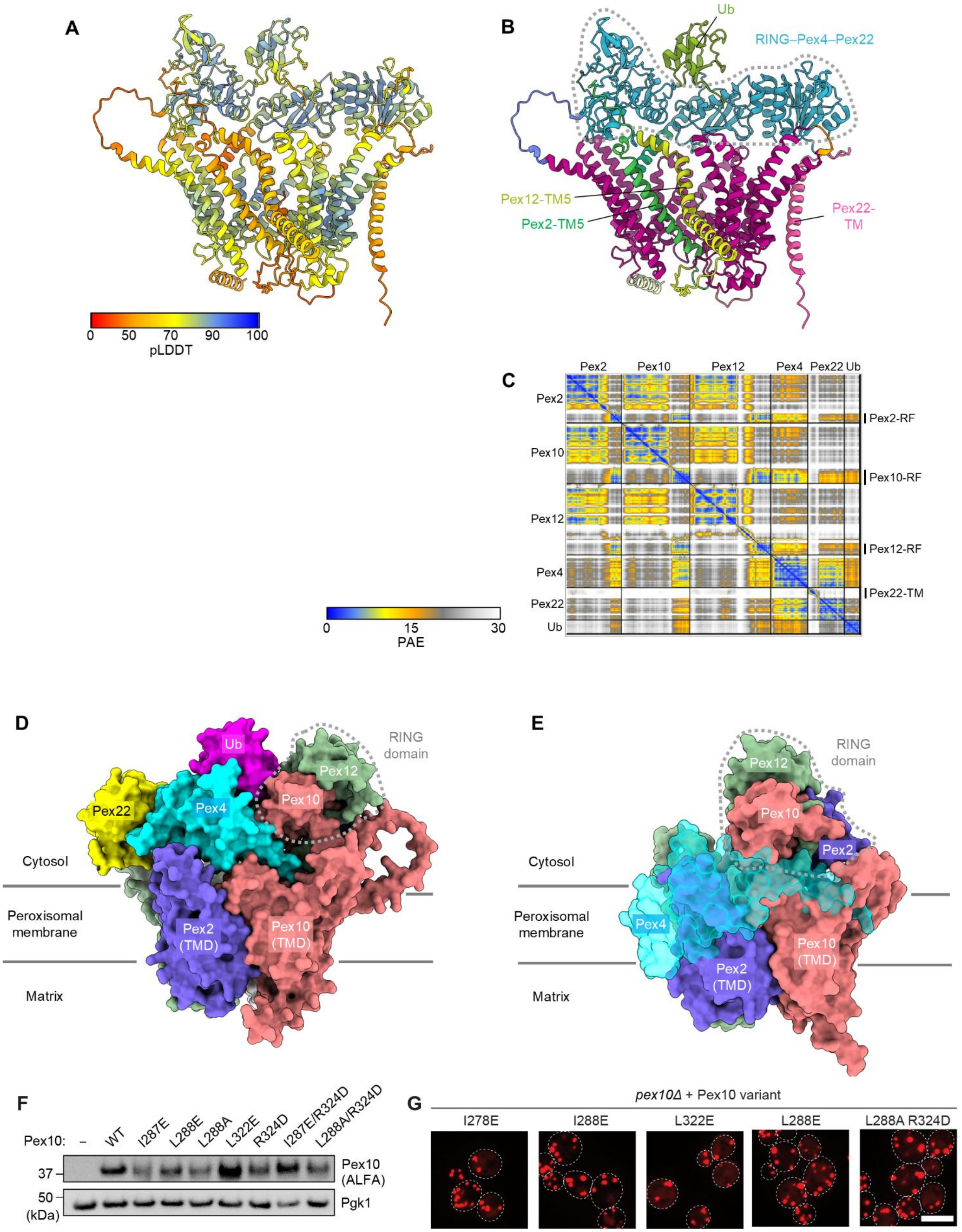

**Supplementary Figure 6. AlphaFold model of the hole-complex containing Pex2-10-12, Pex4, Pex22, and ubiquitin. (see next page for legend)**

**(Legend to Supplementary Figure 6)**
(**A**) pLDDT confidence score map from AlphaFold prediction. (**B**) Domain clustering based on the predicted aligned error (PAE) data visualized in ChimeraX (colored by domain clusters). This analysis identifies the RING domain:Pex4:Pex22 as an interaction group. (**C**) PAE matrix of the prediction. (**D**) As in Fig. 5C, but showing the back side of the model. The fitting highlights excellent placement of Pex4 on the TMDs of Pex2 and Pex10. (**E**) As in Fig. 5E, but showing the back side. Pex4 (semitransparent cyan) was aligned to the closed-state cryo-EM structure of Pex2-10-12–Pex8 according to the canonical E2-RF interaction predicted for Pex4 and Pex10. The model shows major steric clashes between Pex4 and the TMDs of Pex2 and Pex10. (**F**) Immunoblot verifying expression of Pex10 mutants used in Fig. 5G,H. Pex10 contains a C-terminal ALFA-tag. Pgk1, loading control. (**G**) As in Fig. 5H, but showing additional Pex10 mutants. Scale bar, 5 µm.

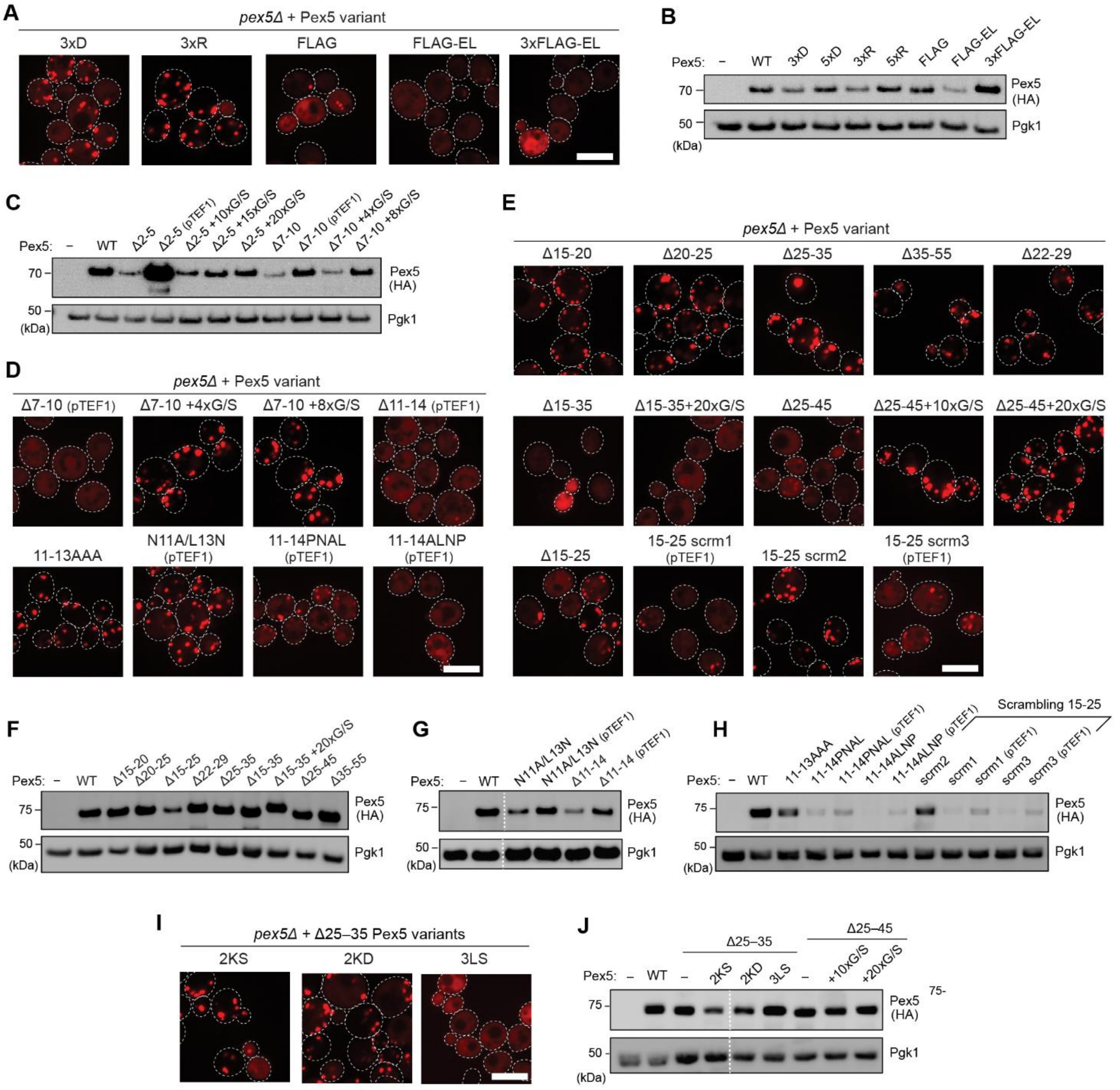

**Supplementary Figure 7. Import activity and expression levels of Pex5 variants.**
As in Fig. 7D, showing results from additional Pex5 variants. Scale bar, 5 µm. (**B**) Immunoblot analysis of Pex5 fused to the indicated N-terminal sequences. All Pex5 constructs carry a C-terminal HA tag. Pgk1, loading control. (**C**) Immunoblot analysis of Pex5 variants with modifications in residues 2–5 or 7–10. (**D**, **E**) Fluorescence microscopy–based mRuby-PTS1 import assay testing Pex5 variants carrying deletions, insertions, or substitution mutations. Scale bar, 5 µm. (**F**–**H**) Immunoblot analysis showing expression levels of Pex5 variants used in D, E, and Fig. 7K–N. (**I**) Fluorescence microscopy–based mRuby-PTS1 import assay testing Δ25–35 Pex5 variants carrying the indicated additional mutations. 2KS = K18S/K24S; 2KD = K18D/K24D; 3LS = L13S/L16S/L36S. Scale bar, 5 µm. (**J**) Immunoblot analysis showing expression levels of Δ25–35 and Δ25–45 Pex5 variants.

**Supplementary Table 1. Cryo-EM data collection, refinement and validation statistics**

| | **Pex2-10-12–Pex8 in the closed state**<br><br>**(EMD-72627; PDB 9Y6Q)** | **Pex2-10-12–Pex8 in the open state**<br><br>**(EMDB-72628; PDB 9Y6R)** |
|---|---|---|
| **Data collection and processing** | | |
| Magnification | 64,000x | 64,000x |
| Voltage (kV) | 300 | 300 |
| Electron exposure ($e^{-}$/Å$^2$) | 50 | 50 |
| Defocus range (μm) | −0.8 to −1.6 | −0.8 to −1.6 |
| Pixel size (Å) | 1.05 | 1.05 |
| Symmetry imposed | *C*1 | *C*1 |
| Initial particle images (no.) | | |
| Final particle images (no.) | 120,987 | 102,420 |
| Map resolution (Å) | 2.94 | 3.05 |
| FSC threshold | 0.143 | 0.143 |
| Map resolution range (Å) | 2.5–7.3<br>(min. to 75$^{th}$ percentile) | 2.6–7.4<br>(min. to 75$^{th}$ percentile) |
| **Refinement** | | |
| Initial model used | AlphaFold model | AlphaFold model |
| Model resolution (Å) | 3.1 | 3.2 |
| FSC threshold | 0.5 | 0.5 |
| Map sharpening *B* factor (Å$^2$) | 129 | 133 |
| Model composition | | |
| Non-hydrogen atoms | 11,617 | 9,824 |
| Protein residues | 1,448 | 1214 |
| Ligands | 7 | 1 |
| *B* factors (Å$^2$) | | |
| Protein | 71.50 | 52.72 |
| Ligand | 109.78 | 31.68 |
| R.m.s. deviations | | |
| Bond lengths (Å) | 0.005 | 0.003 |
| Bond angles (°) | 0.752 | 0.485 |
| **Validation** | | |
| MolProbity score | 1.55 | 1.43 |
| Clashscore | 9.98 | 7.55 |
| Poor rotamers (%) | 0 | 0.09 |
| Ramachandran plot | | |
| Favored (%) | 97.89 | 97.92 |
| Allowed (%) | 2.11 | 2.08 |
| Disallowed (%) | 0 | 0.00 |
| CaBLAM outliers (%) | 1.14 | 0.84 |

**Supplementary Table 2. List of plasmids**

| Plasmid Name | Reference/Notes |
|---|---|
| pYTK1-pYTK96 | Lee et al., 2015 |
| pYTK1-Pex2 (type 3) | This Study |
| pYTK1-Pex10 (type 3) | This Study |
| pYTK1-Pex10-ALFA (type 3) | This Study |
| pYTK1-Pex12-3C-2xSPOT (type 3) | This Study |
| pYTK1-ALFA-Pex8 (type 3) | This Study |
| pYTK1-Pex5 (type 3) | This Study |
| pYTK1-2xHA (type 4a) | This Study |
| pMAL-c2x-MBP-Pex8 | This Study |
| pQE-HisSumo-Pex5-Strep | This Study |
| pYTKe102 [pYTK Hyg/leu2] | For chromosomal integration to the LEU2 locus with hygromycin selection. Assembled from pYTK8, pYTK47, pYTK73, pYTK79, pYTK87, pYTK90, and pYTK93 |
| pYTKe105 [pYTK Kan/HO] | For chromosomal integration to the HO locus with G418 selection. Assembled from pYTK8, pYTK47, pYTK73, pYTK77, pYTK88, pYTK90, and pYTK94 |
| pYTK1-PTS1 (type 4a) | This Study |
| pYTK1-ALFA (type 4a) | This Study |
| pYTK Kan/HO pTDH3-Ub-Y-mRuby2-ePTS1 | This Study |
| pYTK Kan/HO pTDH3-mRuby2-ePTS1 | This Study |
| pYTK95-pTEF1-VioA-pTDH3-VioB | This Study |
| pYTK95 pTEF1-Ub-R-VioE-mVenus-ePTS1 | This Study |
| pYTK Kan/HO pTEF1-VioA-pTDH3-VioB-pTEF1-Ub-R-VioE-mVenus-ePTS1 | This study |
| pYTK96-pGAL1-Pex2, pGAL1-Pex10, pGAL1-Pex12-3C-2xSPOT | This Study |
| pYTK Hyg/leu2 pALD6-Pex5-ALFA | Assembled from pYTK Hyg/leu2, pYTK018, pYTK1-Pex5, pYTKe208, pYTK061 |
| pYTK Hyg/leu2 pALD6-Pex5-2xHA | Assembled from pYTK Hyg/leu2, pYTK018, pYTK1-Pex5, bd07, pYTK061 |
| pYTK Hyg/leu2 pTEF1-Pex5-2xHA | Assembled from pYTK Hyg/leu2, pYTK013, |

| | |
|---|---|
| | pYTK1-Pex5, bd07, pYTK061 |
| pYTK Hyg/leu2 pALD6-ALFA-Pex8 | Assembled from pYTK Hyg/leu2, pYTK018, pYTK1-ALFA-Pex8, pYTK051 |
| pYTK Hyg/leu2 pALD6-Pex10-ALFA | Assembled from pYTK Hyg/leu2, pYTK018, pYTK1-Pex10-ALFA, pYTK051 |
| pYTK Hyg/leu2 pALD6-Pex12-3C-2xSPOT | Assembled from pYTK Hyg/leu2, pYTK018, pYTK1-Pex12-3C-2xSPOT, pYTK051 |
| pWCD1443 | pTEF1-VioA-pTDH3-VioB. (Deloache et al., 2016) |
| pWCD2430 | pTEF1-VioE-YFP-ePTS1. (Deloache et al., 2016) |

**Supplementary Table 3. List of yeast strains**

| Strain ID | Genotype | Reference |
|---|---|---|
| BY4741 | *MATa his3-1, leu2-0, met15-0, ura3-0* | |
| NDY41 | BY4741 *HO::pTDH3-Ub-Y-mRuby2-PTS1-tENO1::KanMX* | this study |
| NDY47 | BY4741 *HO::pTDH3-Ub-Y-mRuby2-PTS1-tENO1::KanMX pex5Δ::LEU2* | this study |
| NDY129 | BY4741 *leu2-0::pALD6-Pex5-ALFA-tENO1::HphMX HO::ScTDH3-Ub-Y-mRuby2-ePTS1-tENO1::KanMX pex5Δ::LEU2* | this study |
| NDY321 | BY4741 *leu2-0::pALD6-Pex5-HA-tENO1-HphMX HO::pTDH3-Ub-Y-mRuby2-PTS1-tENO1::KanMX pex5Δ::LEU2* | this study |
| NDY48 | BY4741 *HO::pTDH3-Ub-Y-mRuby2-PTS1-tENO1::KanMX pex8Δ::LEU2* | this study |
| NDY100 | BY4741 *leu2-0::pALD6-ALFA-pex8-tENO1::HphMX HO::pTDH3-Ub-Y-mRuby2-PTS1-tENO1::KanMX pex8Δ::LEU2* | this study |
| NDY49 | BY4741 *HO::ScTDH3-Ub-Y-mRuby2-ePTS1-tENO1::KanMX pex10Δ::LEU2* | this study |
| NDY103 | BY4741 *leu2-0::pALD6-pex10-ALFA-tENO1::HphMX HO::pTDH3-Ub-Y-mRuby2-PTS1-tENO1::KanMX pex10Δ::LEU2* | this study |
| NDY50 | BY4741 *HO::pTDH3-Ub-Y-mRuby2-PTS1-tENO1::KanMX pex12Δ::LEU2* | this study |
| NDY123 | BY4741 *leu2-0::pALD6-Pex12-3C-2xSPOT-tENO1-HphMX HO::ScTDH3-Ub-Y-mRuby2-ePTS1-tENO1::KanMx pex12Δ::LEU2* | this study |
| yLW006 | BY4741 *ura3-0:pGAL1-Pex2-Pex10-Pex12-3C-2xSPOT::URA3* | This study |

| | | |
|---|---|---|
| yLW019 | BY4741 *HO::pTEF1-VioA-pTDH3-VioB-pTEF1VioE-ePTS1::KanMX* | This study |
| yLW022 | BY4741 *HO::pTDH3-mRuby2-ePTS1::KanMX* | This study |
| yLW025 | BY4741 *HO::pTDH3-mRuby2-ePTS1::KanMX pex5Δ::URA3* | This study |
| yLW050 | BY4741 *HO::pTDH3-mRuby2-ePTS1::KanMX pex8Δ::URA3* | This study |
| yLW051 | BY4741 *HO::pTDH3-mRuby2-ePTS1::KanMX pex10Δ::URA3* | This study |
| yLW052 | BY4741 *HO::pTDH3-mRuby2-ePTS1::KanMX pex12Δ::URA3* | This study |
| yLW340 | BY4741 *leu2-0::ALD6-Pex5-HA WT::HygMX HO::pTDH3-mRuby2-PTS1::KanMX Δpex5::URA3* | This study |
| yLW115 | BY4741 *leu2-0::ALD6-ALFA-Pex8-tENO1::HphMX HO::pTDH3-mRuby2-PTS1::KanMX pex8Δ::URA3* | This study |
| yLW138 | BY4741 *leu2-0::pALD6-Pex10-ALFA::HphMX HO::pTDH3-mRuby2-ePTS1::KanMX pex10Δ::URA3* | This study |
| yLW160 | BY4741 *HO::pTEF1-VioA-pTDH3-VioB-pTEF1-Ub-R-VioE-PTS1::KanMX* | This study |
| yLW170 | BY4741 *HO::pTEF1-VioA-pTDH3-VioB-pTEF1-U-iR-VioE-mVenus-PTS1::KanMX pex12Δ::URA3* | This study |
| yMLT62 | BY4741 *MATa leu2-0::pACT1-GEV::HIS3, rps9Δ, mek1Δ, his3-1, met15-0, ura3-0* | Gift from J. Thorner |

**Supplementary Table 4. List of primers**

| Primer name | Sequence | Notes |
|---|---|---|
| URA3-5F | gggcggattactaccgtt | To confirm plasmid integration at the chromosomal *ura3* locus |
| URA3-5R | gtaatgttatccatgtgggc | To confirm plasmid integration at the chromosomal *ura3* locus |
| URA3-3F | agagcacttgaatccactgc | To confirm plasmid integration at the chromosomal *ura3* locus |
| URA3-3R | gatttggttagattagatatggtttc | To confirm plasmid integration at the chromosomal *ura3* locus |
| LEU2-5F | cataaatacctttcaagc | To confirm plasmid integration at the chromosomal *leu2* locus |
| LEU2-5R | tacaatccttgcccgtgatg | To confirm plasmid integration at the chromosomal *leu2* locus |

| | | |
|---|---|---|
| LEU2-3F | actcgtatcgcatgtcggtg | To confirm plasmid integration at the chromosomal *leu2* locus |
| LEU2-3R | cttcttatgttttacatg | To confirm plasmid integration at the chromosomal *leu2* locus |
| HO-5F | cattcacatcattttcgtggatcc | To confirm plasmid integration at the chromosomal *HO* locus |
| HO-5R | acagcgatggaacttacgg | To confirm plasmid integration at the chromosomal *HO* locus |
| HO-3F | tatcgtgttgcatctgcgg | To confirm plasmid integration at the chromosomal *HO* locus |
| HO-3R | cctttggacttaaaatggcgtg | To confirm plasmid integration at the chromosomal *HO* locus |
| Pex10-YTK1-gDNA-F | gcatcgtctcatcggtctcatatgaagaatgataataagttgcaaaaggaagcacttatg | For amplification of Pex10 CDS from yeast genomic DNA and entry into YTK1 |
| Pex10-YTK1-gDNA-R | atgccgtctcaggtctcaggatctattgccgcaggaccagaatttcctg | For amplification of Pex10 CDS from yeast genomic DNA and entry into YTK1 |
| pex12d-F | aagggaataagcagggaaaataaagcaaaggaaaggaaggtagtcgtcaagtgatgagcttttattcaaacctaccgtctgcaggacaatctagtcgagggctgtggataaccgtagtc | To create pex12Δ yeast strain |
| pex12d-R | atcagattagtagcttcctaatacctgtcacaacttcccatttattcgtgtgtttgttgtgggcgttttttattggtc | To create pex12Δ yeast strain |
| pex12d-conf-F | catcgatgtggacgattac | To confirm deletion of *pex12* by PCR |
| pex12d-conf-R | ctcgagaagattaatcgaatgataatattc | To confirm deletion of *pex12* by PCR |
| pex5d-F | atattgattcaccaaacagttttagttcctattttggatatatatacatcaataaacaatatatcataacacatggacgtaggaagttgctcagctgtggataaccgtagtcg | To create pex5Δ yeast strain |
| pex5d-R | tttataagatttttattgccttatatataggatagcttgtactgctaatgctaatgaatttgggcagtgatgcgagaacataaaattgcggagaaccatagggcgtttttattggtc | To create pex5Δ yeast strain |
| pex8d-5d-conf-F | gctttcatggccctacaa | To confirm deletion of *pex5* and *pex8* by PCR, internal to LEU2 gene |
| pex5d-conf-R | cagcaattatgacctttgaatgc | To confirm deletion of *pex5* by PCR |

| | | |
|---|---|---|
| pex8d-F | ttctaccaggaaaaccgaagaagacaaatgcaaaagggaaatatgaatacgcatgtatgcgcgcaaaaccgcacttacagagggcattaggacattctgtggataaccgtagtcg | To create pex8Δ yeast strain |
| pex8d-R | aggaatataaaaaggcgctactataaagtacttaatgatactataatttagaagattgacttgataagaccgttgtgtaccgggcgtttttattggtc | To create pex8Δ yeast strain |
| pex8d-conf-R | cactttcacttgtatcccagt | To confirm deletion of *pex8* by PCR |
| pex10d-F | attagaaaaataaggtagtggtaataaagcacactacacatcgaagaaagaaacagacactactacatcggtggggggtcatacaatagaggaccaaactgtggataaccgtagtcg | To create pex10Δ yeast strain |
| pex10d-R | acttctcctgctcagacgtctgcgttatttctgaaaaaggcctgtggacaatgctaaaagagtagtcaaattattgattagttcctctattgccgcagggcgtttttattggtc | To create pex10Δ yeast strain |
| pex10d-conf-F | gcgaagtaggtattagccg | To confirm deletion of *pex10* by PCR |
| pex10d-conf-R | gacgtctgcgttatttctgaa | To confirm deletion of *pex10* by PCR |
| pex8-f1-YTK1 | caattctgatatacgtttcgtgcaaagag | To amplify *pex8* from yeast genomic DNA by PCR, removing internal BsaI/BsmBI sites for entry into pYTK1 |
| pex8-r1-YTK1 | ctctttgcacgaaacgtatatcagaattg | To amplify *pex8* from yeast genomic DNA by PCR, removing internal BsaI/BsmBI sites for entry into pYTK1 |
| pex8-f2-YTK1 | atgccgtctcaggtctcaggattactataatttagaagattgacttgataagaccg | To amplify *pex8* from yeast genomic DNA by PCR, removing internal BsaI/BsmBI sites for entry into pYTK1 |
| pex8-r2-YTK1 | gcatcgtctcatcggtctcatatgtttgatcatgacgtcgaatat | To amplify *pex8* from yeast genomic DNA by PCR, removing internal BsaI/BsmBI sites for entry into pYTK1 |
| VioA-B-BsaI-EcoRI | gaaaggtctcaaattccttgccaacaggga g | To generate VioA-VioB with EcoRI/XbaI compatible ends via BsaI endonuclease |
| VioA-B-BsaI-XbaI | gaaaggtctcactagagcttgcaaattaaagcctt | To generate VioA-VioB with EcoRI/XbaI compatible ends via BsaI endonuclease |
| Pex5-YTK1- | gcatcgtctcatcggtctcatatggacgtagg | To amplify *pex5* from yeast genomic |

| | | |
|---|---|---|
| gDNA1-F | aagttgc | DNA by PCR, removing internal BsaI/BsmBI sites for entry into pYTK1 |
| Pex5-YTK1-gDNA1-R | gtaaaataagagtcccaagcatagttg | To amplify *pex5* from yeast genomic DNA by PCR, removing internal BsaI/BsmBI sites for entry into pYTK1 |
| Pex5-YTK1-gDNA2-F | caactatgcttgggactcttattttac | To amplify *pex5* from yeast genomic DNA by PCR, removing internal BsaI/BsmBI sites for entry into pYTK1 |
| Pex5-YTK1-gDNA2-R | gagatccaacgtcaccttttttattattag | To amplify *pex5* from yeast genomic DNA by PCR, removing internal BsaI/BsmBI sites for entry into pYTK1 |
| Pex5-YTK1-gDNA3-F | ctaataataaaaaaggtgacgttggatctc | To amplify *pex5* from yeast genomic DNA by PCR, removing internal BsaI/BsmBI sites for entry into pYTK1 |
| Pex5-YTK1-gDNA3-R | atgccgtctcaggtctcaggatccaaacgaaaattctcctttaaatcttttcag | To amplify *pex5* from yeast genomic DNA by PCR, removing internal BsaI/BsmBI sites for entry into pYTK1 |
| pex8-pMAL-F | ccggaattcgaaaatctttattttcaaggtatgtttgatcatgacgtcg | To amplify *pex8* CDS from pYTK1-AFLFA-Pex8 to clone into pMAL-c2x by digestion with EcoRI/PstI |
| pex8-pMAL-R | aaaactgcagtcagtggtggtggtggtggtgggaccctaatttagaagattgacttgataagaccg | To amplify *pex8* CDS from pYTK1-ALFA-Pex8 to clone into pMAL-c2x by digestion with EcoRI/PstI |
| VioA-B-YTK1-F | gcatcgtctcatcggtctcaaacgccttgccaacagggag | To amplify VioA-VioB cassette with BsmBI cohesive ends from pWCD1443 and insert into YTK1 |
| VioA-B-YTK1-R | atgccgtctcaggtctcacagcagcttgcaaattaaagcctt | To amplify VioA-VioB cassette with BsmBI cohesive ends from pWCD1443 and insert into YTK1 |
| VioE-YTK1-F | gcatcgtctcatcggtctcattctatggagaaccgtgag | To amplify VioE-mVenus-ePTS1 from pWCD2430 and insert into YTK1 |
| VioE-YTK1-R | atgccgtctcaggtctcaggatccctcgagttattacaatttgga | To amplify VioE-mVenus-ePTS1 from pWCD2430 and insert into YTK1 |
